# Colonization of dermal arterioles by *Neisseria meningitidis* provides a safe haven from neutrophils

**DOI:** 10.1101/2021.01.07.425689

**Authors:** Valeria Manriquez, Pierre Nivoit, Tomas Urbina, Hebert Echenique-Rivera, Keira Melican, Patricia Flamant, Taliah Schmitt, Patrick Bruneval, Dorian Obino, Guillaume Duménil

**Author notes:** Current address: Department of Neuroscience, Karolinska Institutet, SE-17177 Stockholm, Sweden. Correspondence: G.D. and D.O.

## Abstract

*Neisseria meningitidis,* a human-specific bacterium, is responsible for meningitis and fatal fulminant systemic disease. Bacteria colonize blood vessels, rapidly causing devastating vascular damage despite a neutrophil-rich inflammatory infiltrate. How this pathogen escapes the neutrophil response is unknown. Using a humanized mouse model, we show that vascular colonization leads to the recruitment of neutrophils, partially reducing bacterial burden and vascular damage. This partial effect is due to the ability of bacteria to indiscriminately colonize capillaries, venules and arterioles, as observed in human samples. In venules, potent neutrophil recruitment allows efficient bacterial phagocytosis. In contrast, in infected capillaries and arterioles adhesion molecules such as E-Selectin are not expressed on the endothelium and intravascular neutrophil recruitment is minimal. These results show that colonization of capillaries and arterioles by *N. meningitidis* create an intravascular niche that preclude the action of neutrophils, resulting in immune escape and subsequent fulminant progression of the infection.

## Introduction

Infections caused by *Neisseria meningitidis* are characterized by an unusually fast progression of the disease as if bacterial proliferation could not be contained by the innate immune system. Although *N. meningitidis* is best known for causing cerebrospinal meningitis, the systemic septic form of the infection is responsible for 90% of the mortality attributed to meningococcal infections^1,2^. In both cases, the infection is characterized by a fulminant evolution and diseases can progress from relatively unspecific “flu-like” symptoms to a life-threatening condition in less than 24 hours^1^. Meningococcal systemic infections are characterized by a typical skin rash called *Purpura* indicating a perturbation of the vascular function^3,4^. Clinical studies have revealed major perturbations in vascular function in infected tissues with congestion, coagulation and the loss of vascular integrity^5,6^. *Neisseria meningitidis* thus rapidly causes major vascular damages leading to a fatal outcome in absence of treatment as if the first lines of immune defence were not sufficient.

How *N. meningitidis* evades innate immunity remains largely unknown. More specifically, an intriguing question that remains is how this bacterium escapes the neutrophil response triggered by the infection. Indeed, both clinical and experimental results point to the recruitment of an inflammatory cellular infiltrate^5,7^ including neutrophils, macrophages and monocytes, which are typically seen around infected blood vessels^5–7^. However, although both neutrophils and monocytic cells frequently contain bacteria^5,7^, this is not sufficient to clear the infection. The recruitment of neutrophils to infected tissues was also observed in a recently developed humanized mouse model in which grafting human skin allows the formation of a network of human dermal vessels that anastomose with the murine dermal vasculature and are perfused with murine blood^8^. In this model, intravenous bacteria exclusively adhere to the human endothelium, progressively proliferate to occupy the entire vessel lumen, and recapitulate the human disease. Taken together, these studies illustrate the ability of the host to initiate an innate immune response upon meningococcal infections, but raise pivotal questions regarding why neutrophils fail to clear these infections.

In this study we hypothesized that the immune escape abilities of *Neisseria meningitidis* are linked to its ability to occupy an intravascular niche. This feature is not unique to *N. meningitidis* but extend to a number of bacterial and viral pathogens that can reach the blood circulation, interact with the endothelium and colonize vessels, including SARS-COV2, as recently described^9–11^. Of note, intravascular colonization by these pathogens generates an atypical situation in terms of neutrophil recruitment. Indeed, the textbook description in which bacteria are found in tissues and neutrophils are recruited to that sites following their rolling on the endothelium of postcapillary venules followed by their extravasation cannot apply for pathogens residing inside blood vessels. Whether the intravascular localization of *Neisseria meningitidis* might account for the incapacity of the innate immune system to clear this infection remains an open question. Therefore, studying the interaction of intravascular bacterial and viral pathogens with endothelial cells is critical to better understand their pathogenesis.

Here, we used the humanized mouse model of meningococcal infection described above, as well as human samples of infected tissues, to explore how neutrophils are recruited to infected vessels and how they target bacteria in this context. We found that *Neisseria meningitidis* escapes this key cellular component of the innate immune response by infecting arterioles and capillaries where neutrophils are not recruited despite intense infection of these vascular beds. *N. meningitidis* thus occupies a specific niche that protects them from the neutrophil response.

## Results

### Dynamics of neutrophil recruitment to sites of infection

As previously shown^8,12^, *Neisseria meningitidis* specifically colonize human vessels within a few hours (Fig. 1a and Supplementary Fig. 1a-b). Intravital time-lapse imaging shows bacteria adhering to the vascular wall (Supplementary Video 1), proliferating in the form of aggregates that become apparent after 2 hours and progressively fuse to finally occupy the vascular space at 6 hours post-infection (*p.i.*). Absence of colonization of mouse vessels confirms the sharp species specificity of the interaction with the endothelium. Efficient heterotypic interactions between murine neutrophils and the human endothelium in our model were confirmed by subcutaneous injection of human TNFα (Supplementary Video 2), as previously described^13^.

**Figure 1.**
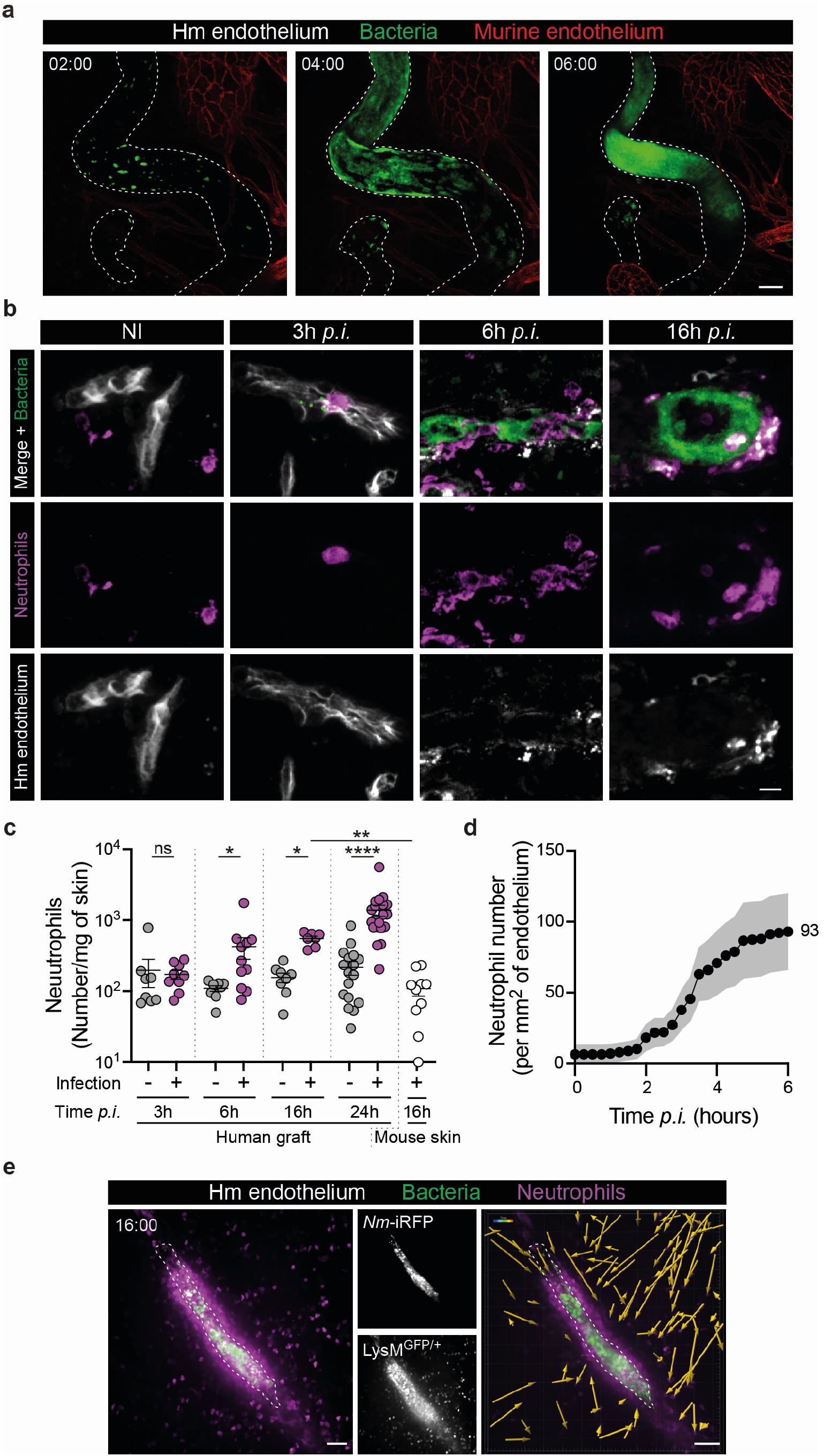
Recruitment of neutrophils to sites of infection. **a,** Representative time sequence of vascular colonization by iRFP-expressing *Neisseria meningitidis* (green). The human and murine endothelia were distinguished using UEA-1 lectin (dashed lines) and mouse-specific anti-CD31 (red) staining, respectively. Time, hh:min. Scale bar, 50 μm. **b,** Representative images of non-infected (NI) and infected human graft tissue sections at indicated times post-infection (*p.i.*) with GFP-expressing *Neisseria meningitidis* (green) and stained for the human endothelium (UEA-1 lectin, grey) and neutrophils (GR-1, magenta). Scale bar, 10 μm. **c,** Flow cytometry quantification of neutrophil numbers in human xenografts of non-infected (−) and infected (+) mice at the indicated times post-infection (*p.i.*). Neutrophil numbers in contralateral mouse skin at 16 hours post-infection is shown on the right. Two-tailed Mann-Whitney test (3h, 6h and 24h time points) and Kruskal-Wallis test with Dunn’s correction for multiple comparisons (16h time point). n≥7 mice per group and time point, in total, pooled from N≥3 independent experiments. **d,** Videos obtained from intravital imaging were used to quantify the numbers of neutrophils per square millimetre of endothelium during the first 6 hours of the infection. Data are shown as the mean±SEM. Quantifications were performed on n=56 vessels, in total, pooled from N=7 infected mice imaged independently. **e,** Intravital imaging (maximum intensity z-projection) of neutrophil (*LysM*^*GFP*/+^, magenta) recruitment at 16 hours post-infection with iRFP-expressing *Neisseria meningitidis* (green) in grafted *Rag*_*2*_^−/−^*γ*_*c*_^−/−^*LysM*^*GFP*/+^ mice. Dashed lines delineate the human endothelium, stained using the UEA-1 lectin. The left panel shows an important swarm of neutrophils around the infected vessel. Time, hh:min. Scale bars, 50 μm. The right panel shows the directed migration of neutrophils towards the infected vessel as revealed by cell tracking over 30 min. Scale bar, 50 μm. ns, not significant; *p<0.05; **p<0.005; ***p<0.0005 and ****p<0.0001.

Neutrophil recruitment to infected vessels was assessed by immunofluorescence of histological samples, flow cytometry, and intravital imaging. Immunohistological staining of infected skin tissues revealed that only a few neutrophils could be found within the infected tissue 3h *p.i.,* with numbers of neutrophils accumulating in the vicinity of infected vessels progressively increasing at 6h *p.i.* onwards (Fig. 1b). Quantitative analysis by flow cytometry of dissociated tissue revealed that neutrophils continued to accumulate up to 24h *p.i.*, whereas contralateral mouse skin tissues did not show this neutrophil accumulation (Fig. 1c and Supplementary Fig. 1c). Intravital imaging of 56 infected vessels confirmed the recruitment of neutrophils during the early phase of the infection (0 to 6h) but only 46% of the infected vessels exhibited at least one neutrophil in their proximity 6h *p.i.*, pointing to a heterogeneous neutrophil response across the infected tissue (Supplementary Fig. 1d). Similarly, the numbers of neutrophils per mm^2^ of endothelium progressively increased and reached the mean value of 93±27 neutrophils 6h *p.i.* (Fig. 1d).

Reminiscent of human cases^7^, at 16h *p.i.*, the recruitment of *LysM*^*GFP*^-labelled neutrophils visualized by intravital imaging was massive with certain infected vessels being surrounded by large numbers of neutrophils, forming a sheath around the vessel (Fig. 1e and Supplementary Video 3). Monocytes also express the *LysM*^*GFP*^ reporter but were easily distinguished from neutrophils by their low level of fluorescence^14^. Single cell tracking showed that neutrophils converge from the parenchyma towards the infected vessel (Fig. 1e and Supplementary Video 3). Together these observations demonstrate that neutrophils progressively accumulate in the vicinity of certain infected vessels following *N. meningitidis* vascular colonization, starting at 3 hours post-infection and reaching large amounts at 16-24h *p.i.*, as observed in human cases.

### Intravascular colonization determines the nature and timing of neutrophil recruitment

We next investigated the bacterial signals involved in neutrophil recruitment. To determine whether neutrophil recruitment relied on vascular colonization by *N. meningitidis*, we studied mutants that have altered ability to interact along the vascular wall. Grafted mice were first infected with two isogenic bacterial strains, *pilC1*, a piliated but non-adherent mutant, and *pilD*, a mutant that does not express any pili on its surface^15^ (Fig. 2a). Both strains cannot adhere to the endothelium and thus cannot colonize vessels. As expected^8^, while mice were inoculated with similar amounts of bacteria (Fig. 2b), the numbers of adherent bacteria 24h *p.i.* were strongly decreased when mice were infected with the mutant bacterial strains (Fig. 2c). We found that neutrophil recruitment 24h after infection was tightly dependent on bacterial adhesion since their numbers barely increased when mice were infected with either of the mutant bacterial strains (Fig. 2d).

**Figure 2.**
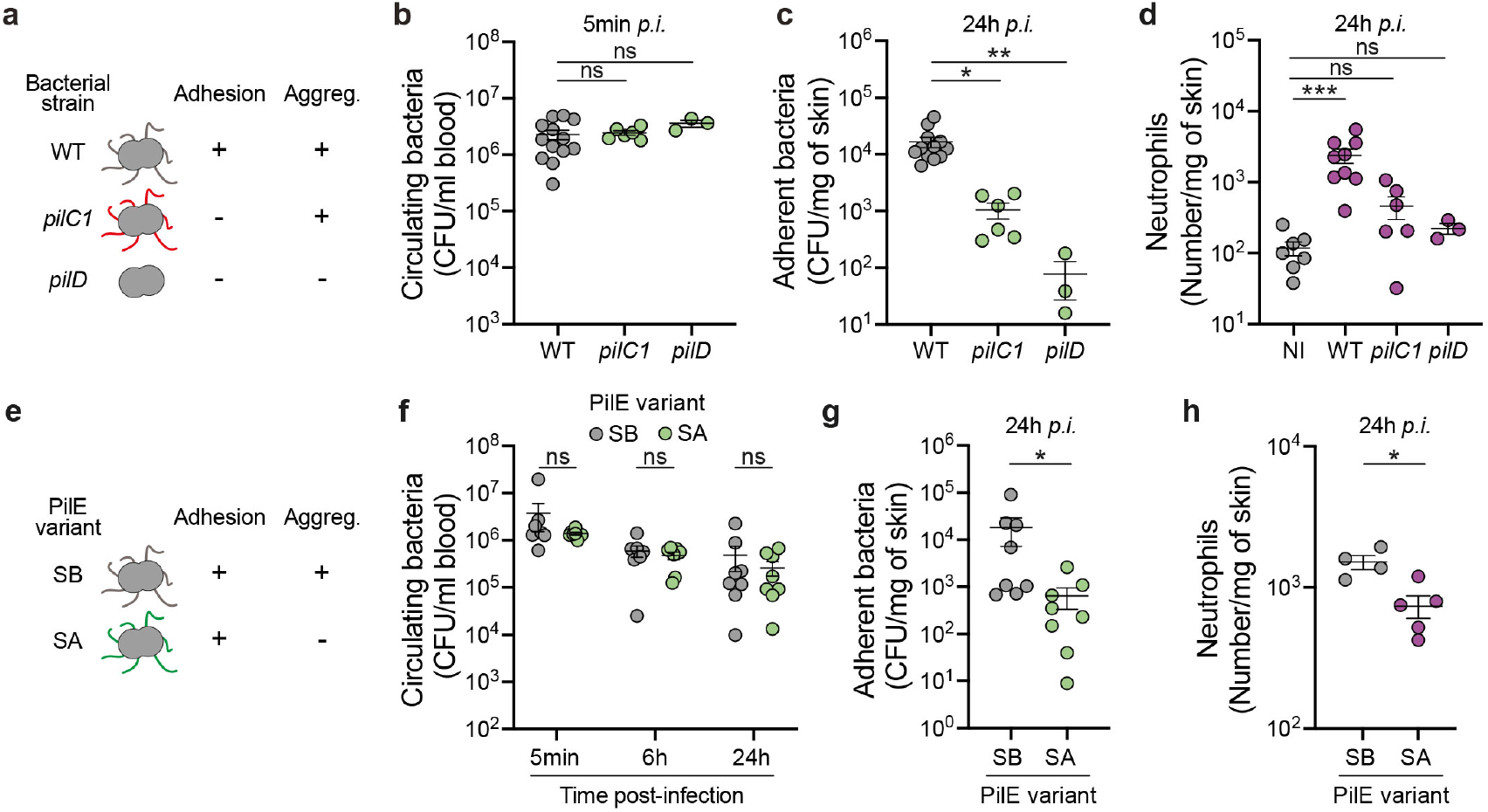
Neutrophil recruitment to infected vessels relies on type IV pili-mediated bacterial adhesion and aggregation. **a,** Schematic representation of the different bacterial strains used to assess the impact of bacterial adhesion on neutrophil recruitment. **b,** Bacterial colony forming unit (CFU) counts from blood (circulating bacteria) of mice infected for 5 minutes with either WT, *pilC1* (piliated but non-adherent) or *pilD* (non-piliated and non-adherent) bacterial strains. **c,** Bacterial colony forming unit (CFU) counts from dissociated human xenografts (adherent bacteria) collected from mice infected for 24 hours with either WT, *pilC1* or *pilD* bacterial strains. **d,** Flow cytometry quantification of neutrophil numbers in human xenografts harvested from non-infected mice and mice infected for 24 hours with either WT, *pilC1* or *pilD* bacterial strains. **a-c,** Kruskal-Wallis test with Dunn’s correction for multiple comparisons. n≥6 mice per group (except for *pilD* n=3), in total, pooled from N=3 independent experiments (except for *pilD* N=1). **e,** Schematic representation of the different bacterial strains used to assess the impact of bacterial aggregation on neutrophil recruitment. **f,** Bacterial colony forming unit (CFU) counts from blood (circulating bacteria) of mice infected for the indicated times with either the wild type strain (SB) or a non-aggregative strain (SA). Two-tailed Mann-Whitney test per time point. n≥7 mice per group, in total, pooled from N≥3 independent experiments. **g,** Bacterial colony forming unit (CFU) counts from dissociated human xenografts (adherent bacteria) collected from mice infected for 24 hours with either SB or SA pilin variant-expressing bacteria. Two-tailed Mann-Whitney test. n≥8 mice per group, in total, pooled from N≥3 independent experiments. **h,** Flow cytometry quantification of neutrophil numbers in human xenografts harvested from mice infected for 24 hours with either SB or SA pilin variant-expressing bacteria. Two-tailed Mann-Whitney test. n≥4 mice per group, in total, pooled from N=2 independent experiments. ns, not significant; *p<0.05; **p<0.005; ***p<0.0005 and ****p<0.0001.

Type IV pili also allow bacteria to form three-dimensional viscous aggregates containing thousands of bacteria and participating in vessel occlusion^12^. To test the impact of these bacterial aggregates inside blood vessels on neutrophil recruitment, a strain expressing a sequence variant (SA) of the major pilin unable to generate auto-aggregation while still adhering to endothelial cells^16^ was compared to the wild type sequence variant (SB) (Fig. 2e). While both strains showed the same ability to survive inside the circulation, the SA variant showed lower amounts of bacteria colonizing blood vessels as expected (Fig. 2f-g). The amounts of neutrophils recruited by the non-aggregative strain was reduced (Fig. 2h), showing that the formation of bacterial aggregates enhanced neutrophil recruitment. These data show that bacterial adhesion, amplified by bacterial auto-aggregation, both mediated by type IV pili, is essential to initiate a cascade eventually leading to neutrophil recruitment.

### Neutrophils only partially control the number of colonizing bacteria and subsequent vessel damages

To address the role of recruited neutrophils during meningococcal disease, we depleted these cells and evaluated the effect on infection. Grafted mice were pre-treated with the monocytes and neutrophil-depleting antibody directed against GR-1 (clone RB6-8C5), 24h prior to infection (Fig. 3a). This treatment led to an efficient depletion of circulating neutrophils when compared to the mice that received the isotype control antibody (Fig. 3b). Absence of neutrophils in the circulation did not affect the number of non-adherent circulating bacteria after intravenous injection at any of the tested times post-infection (Fig. 3c). As expected, the depletion effectively prevented the neutrophil recruitment to infected human tissues (Fig. 3d). Importantly, the number of adherent bacteria at 24h *p.i.* was strongly increased when compared to the control condition (Fig. 3e). Nearly identical results were obtained when the mice were treated with the neutrophil-specific depletion antibody clone 1A8, against Ly-6G, supporting that monocytes likely do not further contribute to bacterial clearance (Supplementary Fig. 2a-d). These results indicate that neutrophils are the primary cell type controlling the number of bacteria adhering and proliferating along the endothelium, but are unable to completely resolve the infection within this time frame.

**Figure 3.**
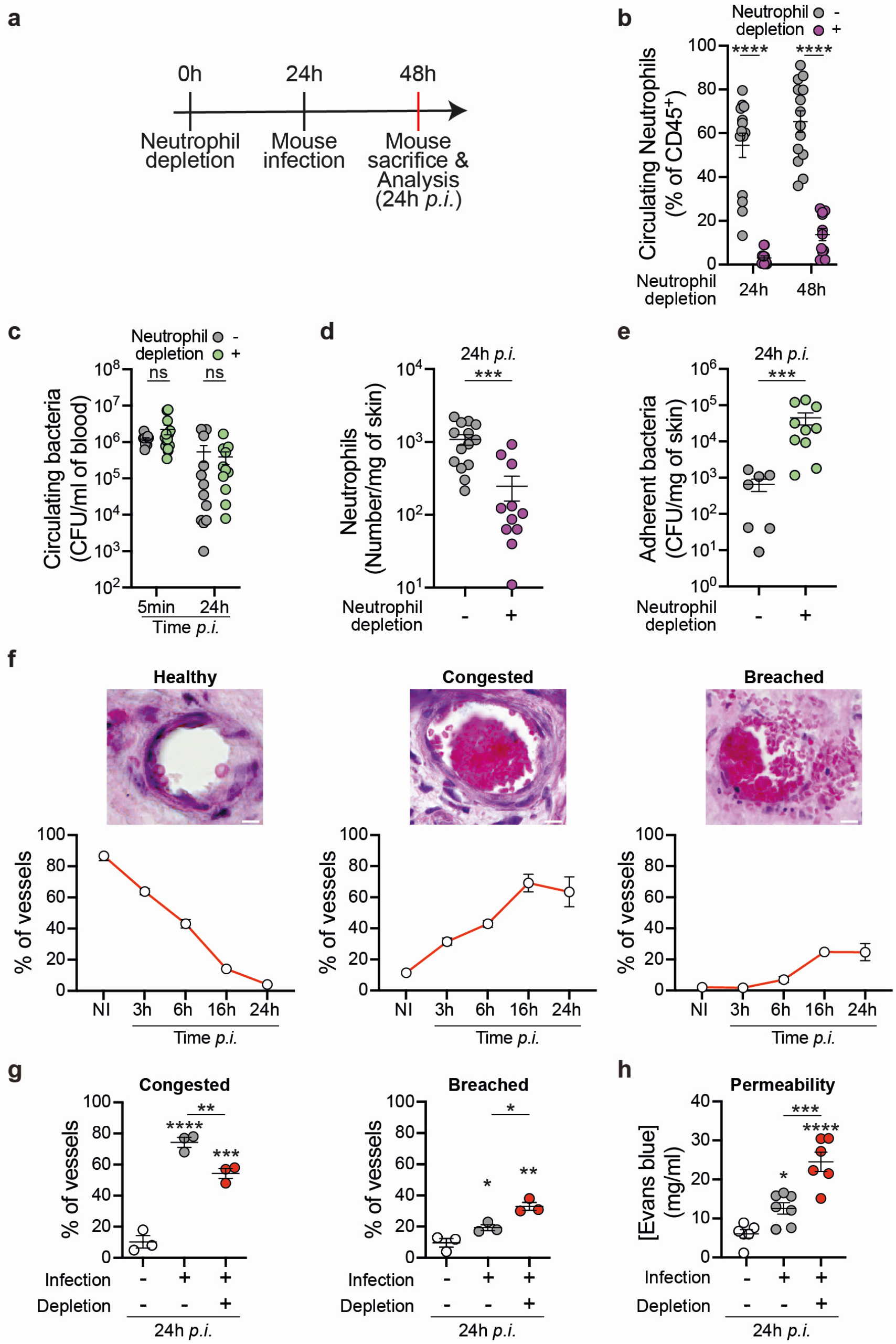
Neutrophils limit vascular colonization by meningococci and vessel damages. **a,** Schematic of the experimental approach used: neutrophil depletion was achieved by intravenous injection of the neutrophil-depleting antibody (anti-GR-1, clone RB6-8C5) 24h prior to mouse infection. Mice were sacrificed 24h post-infection and analyses were carried out. **b,** Numbers of blood circulating neutrophils in non-infected mice pre-treated with either the isotype control antibody (−) or the neutrophil-depleting antibody (+). **c,** Bacterial colony forming unit (CFU) counts from blood (circulating bacteria) of mice pre-treated with either the isotype control antibody (−) or the neutrophil-depleting antibody (+) and infected for 5 minutes or 24 hours. **d,** Neutrophil numbers in human xenografts of mice pre-treated with either the isotype control antibody (−) or the neutrophil-depleting antibody (+) and infected for 24 hours. **b-d,** Two-tailed Mann-Whitney test per time point. n≥11 mice per group, in total, pooled from N=4 independent experiments. **e,** Bacterial colony forming unit (CFU) counts from dissociated human xenografts (adherent bacteria) collected from mice pre-treated with either the isotype control antibody (−) or the neutrophil-depleting antibody (+) and infected for 24 hours. Two-tailed Mann-Whitney test. n≥7 mice per group, in total, pooled from N=3 independent experiments. **f,** Representative images of Haematoxylin and Eosin (H&E) stained histological sections of infected human xenografts and quantification of vascular damage in the infected vessels. Vessels were classified as healthy, congested (intraluminal accumulation of red blood cells with abnormal morphology and fibrin clot) or breached (presence of red blood cells out of the vessels). Quantifications were performed on n≥500 vessels, in total, pooled from N≥3 mice per time point. Scale bar, 10 μm. **g,** Quantification of vascular damage upon mouse infection and neutrophil depletion using the GR-1 antibody (clone RB6-8C5). One-way ANOVA with Holm-Sidak’s correction for multiple comparisons. Quantifications were performed on n≥200 vessels, in total, pooled from N≥3 mice per group. **h,** Quantification of vascular permeability upon mouse infection and neutrophil depletion with the anti-Ly-6G antibody (clone 1A8) using Evans blue. One-way ANOVA with Holm-Sidak’s correction for multiple comparisons. n≥6 mice per group, in total, pooled from N=4 independent experiments. ns, not significant; *p<0.05; **p<0.005; ***p<0.0005 and ****p<0.0001.

As pathological effects of *Nm* infections are linked to altered vascular function, we also measured the potential contribution of neutrophils to vascular damage. Indeed, neutrophil have been reported to induce vascular damage in other systems^17^, and infection by *Neisseria meningitidis* in humans have been shown to involve vascular damage^7,8^. Kinetics of vascular damage after *N. meningitidis* infection were first determined using histology and intravital imaging to compare with neutrophil recruitment kinetics. Over 500 blood vessels on histological slices were analysed for each time point and classified as healthy, congested (with accumulation of red blood cells), or breached (with perivascular extravasation of red blood cells). Representative images and quantitative results of each vascular condition are shown in Figure 3f. The percentage of healthy vessels gradually declined down to only 4% 24h *p.i.*. Conversely, congestion and vascular rupture progressively increased with time. Interestingly, a significant increase in vascular congestion was observed as early as 3h *p.i.* and thus before recruitment of neutrophils. The percentage of congested vessels continued to increase to reach a maximum percentage of 70% at 16h *p.i.*, which remained stable until 24h *p.i.* A similar situation occurred with vascular rupture, with a slight increase at 6h *p.i.* from 2% to 7% that progressively increased, reaching 25% at 16h *p.i.*.

Neutrophil depletion led to a decrease in the number of congested vessels and a concurrent increase in the occurrence of breached vessels with released red blood cells (Fig. 3g and Supplementary Fig. 2e). To confirm and extend these results, vessel permeability to serum content was then measured using Evans Blue^18^. This dye was injected intravenously and 10 min post-injection, the circulating dye was removed from blood vessels by a myocardial perfusion of heparin-containing buffer. As expected, a 24h infection led to an increase of vascular permeability evidenced by high Evans blue accumulation in the tissue. In the absence of neutrophils, vascular leakage due to the infection was further increased (Fig. 3h). Altogether these data are in favour of a protective role for neutrophils during meningococcal infection. After being recruited, neutrophils incompletely control the number of bacteria in the infected vessels and thus partially reduce the vascular damage induced by the bacteria upon vascular colonization.

### Ability of bacteria to colonize distinct vascular beds determine the level of neutrophil recruitment

Our results at this stage reveal a paradox, neutrophils are recruited upon vascular colonization and provide protection but fail to prevent the initiation and progression of the infection. A possible explanation of this incomplete neutrophil response is that the recruitment is insufficient in terms of number. This hypothesis is supported by the relatively low overall numbers of neutrophils (93±27 neutrophils.mm^−2^ at 6h *p.i*.) that are observed following *N. meningitidis* infection. As a comparison, TNFα stimulation or similar inflammatory situations can lead to about 5-fold this value^19^. We thus hypothesized that the unusual intraluminal location of the bacteria, rather than the bacterium itself, is responsible for this moderate recruitment. To test this hypothesis, bacteria were injected directly into the xenograft intradermally and the inflammatory infiltrate evaluated. The amounts of injected bacteria were adjusted to reach the amount found following intravascular infection (Fig. 4a). In these conditions, histological analysis of the human skin graft 3h post-intradermal infection showed a considerable infiltration of inflammatory cells in contrast to intravascular infection at the same time point (Fig. 4b). This was confirmed by flow cytometry analysis that highlighted a 10-fold increase in neutrophil numbers as early as 3h post-intradermal infection (Fig. 4c). Even at 6h post intravascular infection such numbers were not reached (Fig. 1c). These results point to an unexpectedly slow kinetics of neutrophil recruitment during vascular colonization and a particular impact of the vessel intraluminal location of the bacteria during infection.

**Figure 4.**
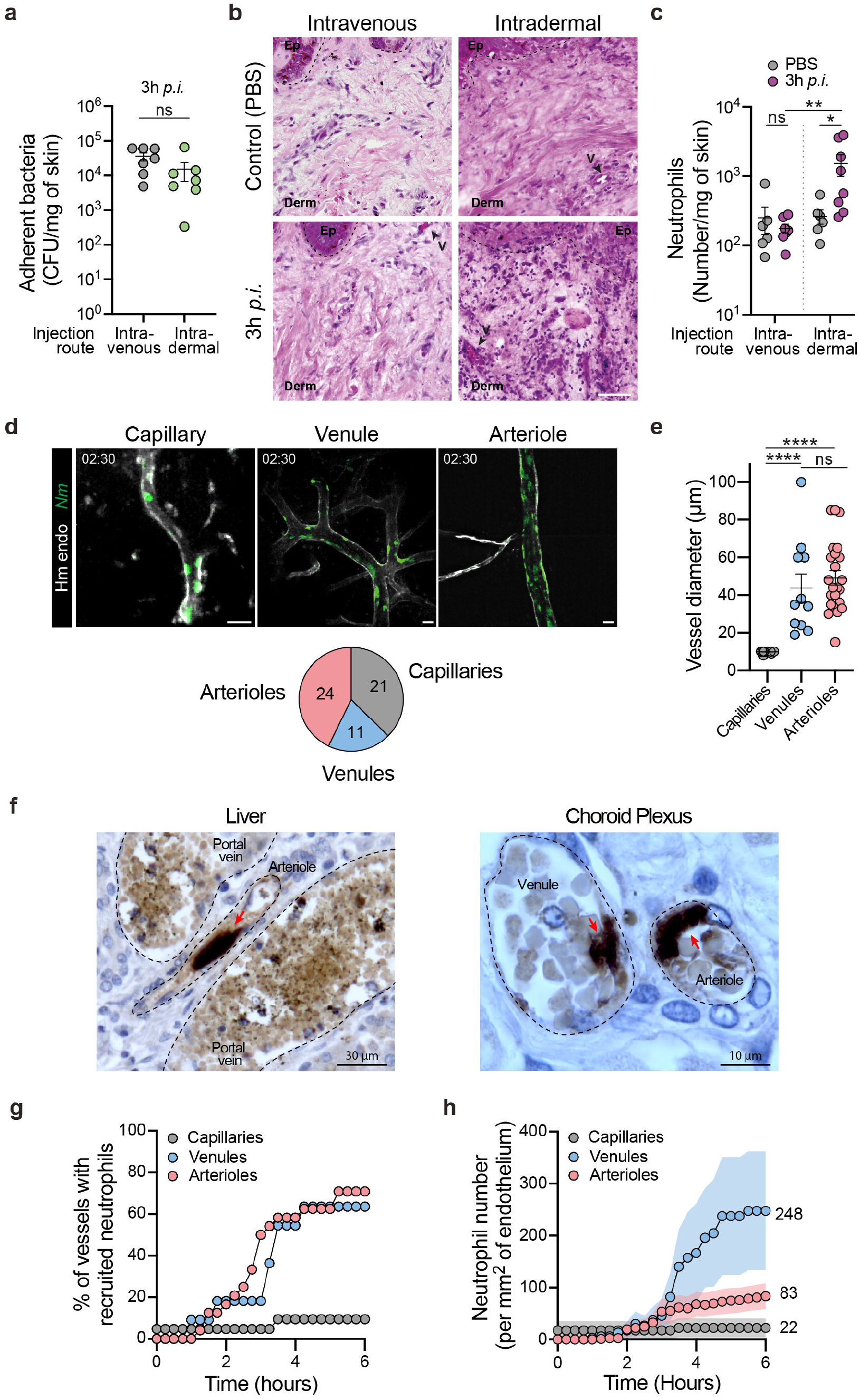
Different types of vessels are infected with different levels of neutrophil recruitment. **a,** Bacterial colony forming unit (CFU) counts from dissociated human xenografts (adherent bacteria) collected from either intravenously or intradermally infected mice for 3 hours. Mann-Whitney test. n=7 mice, in total, pooled from N=3 independent experiments. **b,** Representative images of Haematoxylin and Eosin stain of human xenografts harvested from mice 3 hours following either an intravenous or an intradermal injection of PBS (control) or *Neisseria meningitidis* (3h *p.i.*). Scale bar, 50 μm. The dashed lines delineate the epidermis/dermis border. Arrowheads indicate blood vessels. V, vessel. **c,** Neutrophil numbers in human xenografts of control and infected (3h *p.i.*) mice by either intravenous or intradermal injection. Kruskal-Wallis test with Dunn’s correction for multiple comparisons. n≥6 mice per group, in total, pooled from N=3 independent experiments. **d,** Representative images of vascular colonization of the different vascular beds: capillaries, venules, and arterioles, 2h30 post-infection. The pie chart highlights the proportion and number of each vessel type. Scale bar, 20 μm. **e,** Quantification of capillary, venule, and arteriole diameter. Kruskal-Wallis test with Dunn’s correction for multiple comparisons. **f,** Immunohistochemistry analysis of a case of meningococcal septic shock showing vascular colonization by *N. meningitidis* in both venules and arterioles in the liver (left) and the choroid plexus (right). Bacteria were labelled using a polyclonal antibody directed against the strain cultured from the blood of this patient, as previously described^23^. **g,** Percentage of vessels recruiting at least one neutrophil following infection according to the different vascular beds. **h,** Numbers of neutrophils per square millimetre of endothelium according to the different vascular beds during the first 6 hours of the infection. **d-e and g-h,** Quantifications were performed on n=56 vessels (21 capillaries, 11 venules and 24 arterioles), in total, pooled from N=7 infected mice imaged independently. ns, not significant; *p<0.05; **p<0.005; ***p<0.0005 and ****p<0.0001.

It is important to note that numbers given above represent average values throughout the tissue that could be highly heterogeneous and mask specific locations. We thus analysed in depth whether bacteria equally infected distinct types of vessels. For this, infected vessels were categorized into three classes: capillaries, venules and arterioles. The distinction between venules and arterioles was based on a combination of parameters: morphology, intensity and direction of blood flow as well as an arteriole-specific staining approach, as previously described^20^ (Supplementary Fig. 3a-b). Capillaries were characterized by a luminal diameter below or equal to 10 μm. Remarkably, this classification revealed that *N. meningitidis* could infect all vessel types (Fig. 4d). Observed infected venules and arterioles had a similar diameter, 43.73±7.52 μm and 49.13±3.77 μm, respectively (Fig. 4e). In order to validate these observations made in an experimental mouse model, evidence of colonization of venules and arterioles in a human case were sought. *Post-mortem* samples from a case of *purpura fulminans* were immuno-stained to visualize bacteria and tissue organization. Evidence of colonization of both venules and arterioles could be found in different tissues, such as in liver and choroid plexus, in which aggregates of bacteria were associated to the endothelium (Fig. 4f, red arrows). *N. meningitidis* is thus able to colonize all types of vascular beds both in an experimental system and during human infections.

Neutrophil rolling was shown to preferentially occur on the venular endothelium but to a much lesser extent on arterioles and capillaries^21^. We therefore hypothesized that they might not be efficiently recruited to bacteria colonies localized in arterioles and capillaries. To test this hypothesis, the recruitment of neutrophils to the different vessel beds was evaluated. Neutrophils were rarely recruited to infected capillaries (Fig. 4g). The kinetics and percentage of infected vessels recruiting at least a single neutrophil was similar between venules and arterioles showing that some level of recruitment occurred in both of these vessel types (Fig. 4g). Importantly, however, the number of neutrophils recruited was much higher in venules than in arterioles, with 248±115 *versus* 83±25 neutrophils per mm^2^ of endothelium, respectively at 6h *p.i.* (Fig. 4h). The number of neutrophils recruited to capillaries was even lower than in arterioles (22±18 neutrophils per mm^2^ of endothelium). The ability of meningococci to colonize arterioles could thus provide an environment with lower numbers of neutrophils and thus an edge over the innate immune system.

### Neutrophils recruited to venules efficiently phagocytose bacteria

Results described above show a preferential recruitment of neutrophils to infected venules. We thus wondered whether neutrophils could have efficient bacterial killing properties in this context. The recruitment of neutrophils to infected venules was therefore explored in further detail in terms of location, kinetics and phagocytic activity.

Dynamic intravital imaging of infected venules revealed the expected progressive accumulation of bacteria along the vascular walls (Fig. 5a). Quantitative analysis of 11 venules revealed that after 4 hours of infection, 55% of infected venules displayed neutrophil recruitment inside the vascular lumen (Fig. 5b). Rarely, neutrophils could be found in a perivascular location (1/11 venules). Numbers of intraluminal neutrophils reached 216±91 neutrophils per mm^2^ of endothelium after 6 hours of infection (Fig. 5c). Neutrophils on the endothelium were highly motile and efficiently explored the infected endothelium by crawling on its surface (Supplementary Video 4). On frequent occasions neutrophils crawled in the direction of adherent bacterial aggregates. Neutrophils could efficiently internalize the adherent bacteria and thus detach them from the cellular surface (Fig. 5d and Supplementary Video 4). During long-term observations, bacterial aggregates initially adhering to the vascular wall could no longer be found following the arrival of neutrophils, demonstrating their efficiency in this context (Fig. 5a, red arrows and Supplementary Video 4). This implies that neutrophils have the ability to detach bacteria tightly bound to the endothelial surface. Collectively, these experimental and clinical observations suggest that meningococci can adhere to venules and that neutrophils are readily recruited to these sites where they can efficiently phagocytose bacteria adhering to the endothelial wall.

**Figure 5.**
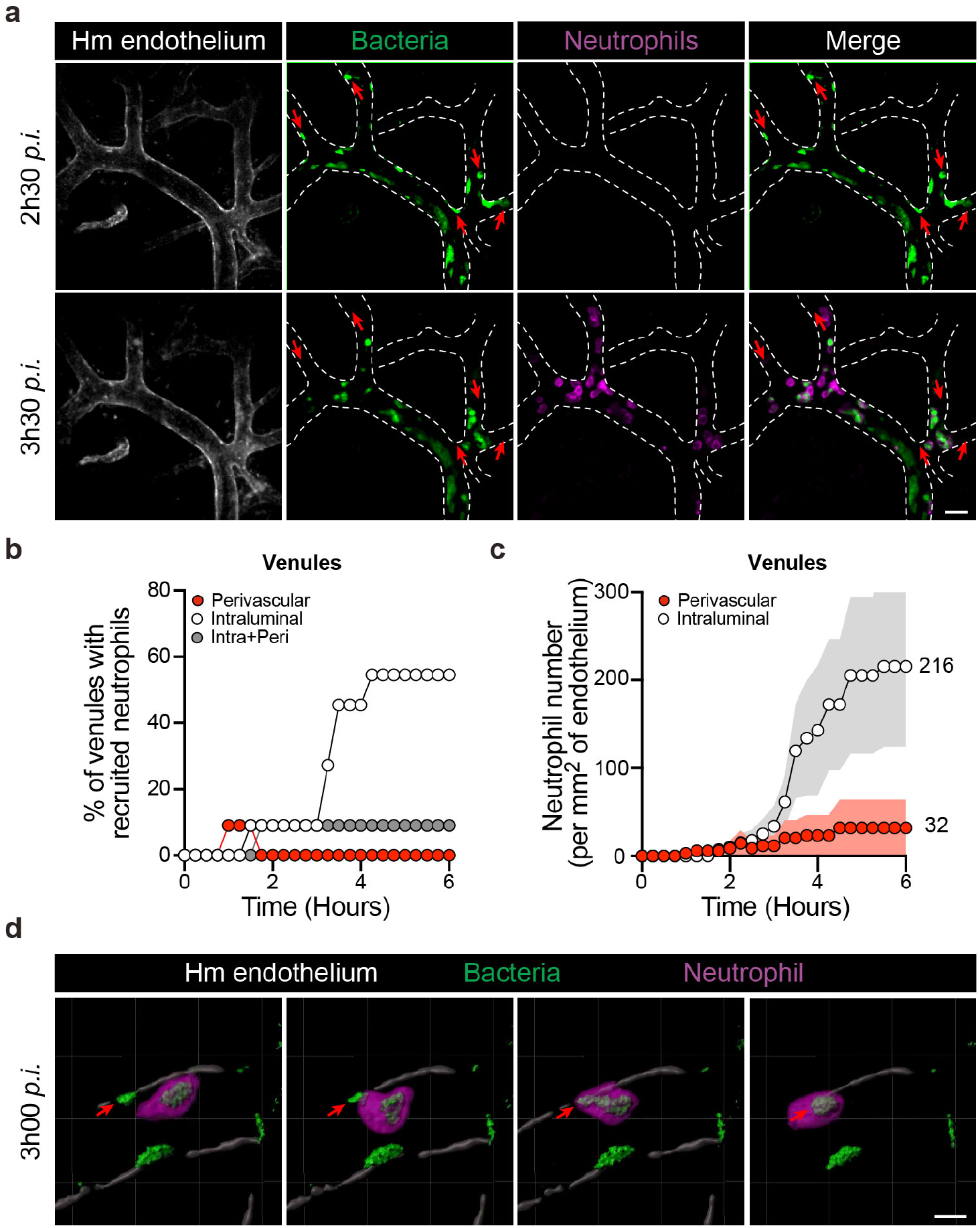
Vascular colonization of venules leads to efficient neutrophil recruitment and function. **a,** Intravital imaging of neutrophil (Ly-6G, magenta) recruitment to infected venules with iRFP-expressing *Neisseria meningitidis* (green). The human vessels are shown in grey (UEA-1 lectin) and dashed lines. Scale bars, 30 μm. **b,** Percentage of venules recruiting only intraluminal (white), only perivascular (red) or both intraluminal and perivascular (grey) neutrophils. **c,** Numbers of intraluminal and perivascular neutrophils per square millimetre of venular endothelium during the first 6 hours of the infection. **b-c,** Quantifications were performed on n=11 vessels, in total, pooled from N=7 infected mice imaged independently. **d,**3D-rendering of intravital imaging showing neutrophil (Ly-6G, magenta) migration toward and engulfing an adherent bacterial aggregate within a venule 3h post-infection with iRFP-expressing *Neisseria meningitidis* (green). The human venule is shown in grey (UEA-1 lectin). Scale bar, 10 μm.

Our results show that in the context of arterioles only few neutrophils were recruited following infection, leading to the hypothesis that the neutrophil response would be inefficient in this vessel type. The recruitment of neutrophils to arterioles was then examined in detail. Intravital imaging revealed that the few neutrophils recruited to arterioles were either located in an intra- or extra-luminal location, occasionally both (Fig. 6a). Neutrophil location was quantified in 24 arterioles over time, revealing that intraluminal and extraluminal or perivascular locations were represented in similarly proportions (Fig. 6b). During the course of the infection, the recruitment of both intraluminal and perivascular neutrophils progressively increased over time with similar kinetics and similar low numbers per surface of endothelium (Fig. 6c). The few intraluminal neutrophils were frequently found to contain large amounts of bacteria (Fig. 6a, red arrows). In contrast, perivascular neutrophils were never observed to contain bacteria and even though neutrophils were often closely wrapped around to the vascular wall (Supplementary Video 5). No evidence of perivascular neutrophils translocating into infected arterioles was observed. Following 4-6 hours of infection, vessels were clogged with bacteria and intraluminal neutrophils progressively showed reduced motility (Fig. 6d and Supplementary Video 6). These results show that when arterioles are infected, only limited amounts of neutrophils are recruited and furthermore their efficiency is limited. The few intravascular neutrophils are rapidly overwhelmed with bacteria filling the vessel lumen. Extravascular neutrophils fail to enter vessels and access bacteria.

**Figure 6.**
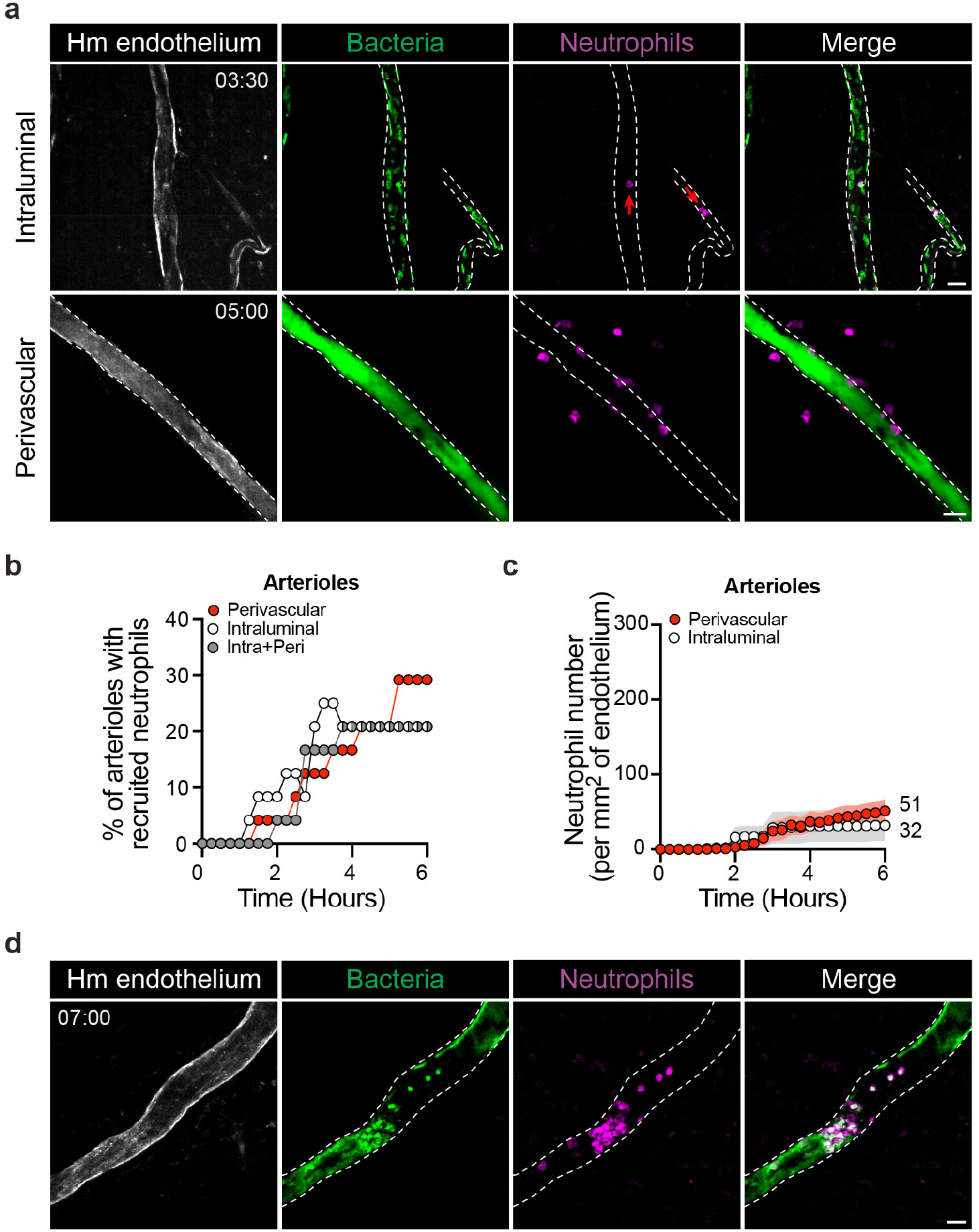
Vascular colonization of arterioles leads to limited neutrophil recruitment and efficiency. **a,** Intravital imaging of intraluminal (top panel) or perivascular (bottom panel) recruitment of neutrophils (Ly-6G, magenta) to arterioles 3-5 hours post-infection with iRFP-expressing *Neisseria meningitidis* (green). The human vessels are shown in grey (UEA-1 lectin) and dashed lines. Time, hh:min. Scale bar, 30 μm. **b,** Percentage of arterioles recruiting only intraluminal (white), only perivascular (red) or both intraluminal and perivascular (grey) neutrophils. **c,** Numbers of intraluminal and perivascular neutrophils per square millimetre of arteriolar endothelium during the first 6 hours of the infection. **b-c,** Quantifications were performed on n=24 vessels, in total, pooled from N=7 infected mice imaged independently. **d,** Intravital imaging of neutrophil (Ly-6G, magenta) entrapped within bacterial aggregates within an infected arteriole 7h post-infection with iRFP-expressing *Neisseria meningitidis* (green). The human vessels are shown in grey (UEA-1 lectin) and dashed lines. Scale bar, 30 μm.

Therefore, the ability of meningococci to adhere to different vascular beds and more specifically arterioles and capillaries allow them to rapidly colonize these vessels before an efficient innate immune response can be mounted.

### Despite intense colonization endothelia lining capillaries and arterioles fail to upregulate inflammatory adhesion molecules

The above results raised the question of why neutrophils were efficiently recruited to venules but not to capillaries and arterioles despite large accumulation of bacteria in all these vessel types. During infections caused by other pathogens, occurring inside tissues, neutrophils specifically exit the circulation through venules to reach the infection site. In this case, the specificity for venules is linked to the expression of a set of adhesion receptors on the endothelium surface, which are triggered by inflammatory signals coming from the infection site^21^. Although the intravascular location of the *N. meningitidis* infection generates a different situation, the difference in neutrophil response could be due to a differential expression of adhesion receptors. We first determined the consequence of meningococcal infection on the expression of a selection of neutrophil adhesion receptors on human umbilical vein endothelial cells *in vitro*. While ICAM-1 and VCAM-1 cell surface levels barely increase 5h after infection, the expression of E-Selectin (CD62E) was strongly induced by the infection (Fig. 7a), as previously reported^22^. Expression of E-Selectin was highest at 5h post-infection but was clearly detectable as early as 2h post-infection. This kinetics of expression being compatible with the kinetics of neutrophils recruitment observed during infection *in vivo,* we then determined the expression of E-Selectin in the humanized mouse model of *N. meningitidis* infection. Low concentrations of fluorescently labelled anti-human CD62E monoclonal antibody were introduced in the circulation of infected animals and accumulation of signal along infected vessel walls followed. Infection in the context of capillaries and arterioles did not lead to any specific signal (Fig. 7b), where only a low level and constant background signal could be seen. In contrast, infected venules showed a gradual accumulation of E-Selectin signal during the course of the infection (Fig. 7b). Interestingly, the signal was heterogeneous along the human endothelium surface and strongly colocalized with bacterial colonies (Fig. 7c). Signals leading to E-selectin expression thus had a local component linked to bacterial adhesion. Together, these results show that despite massive infection and in contrast to the venular endothelium, endothelia from arterioles and capillaries fail to express E-selectin on their surface upon infection, thus preferentially targeting neutrophil recruitment towards infected venules over arterioles and capillaries.

**Figure 7.**
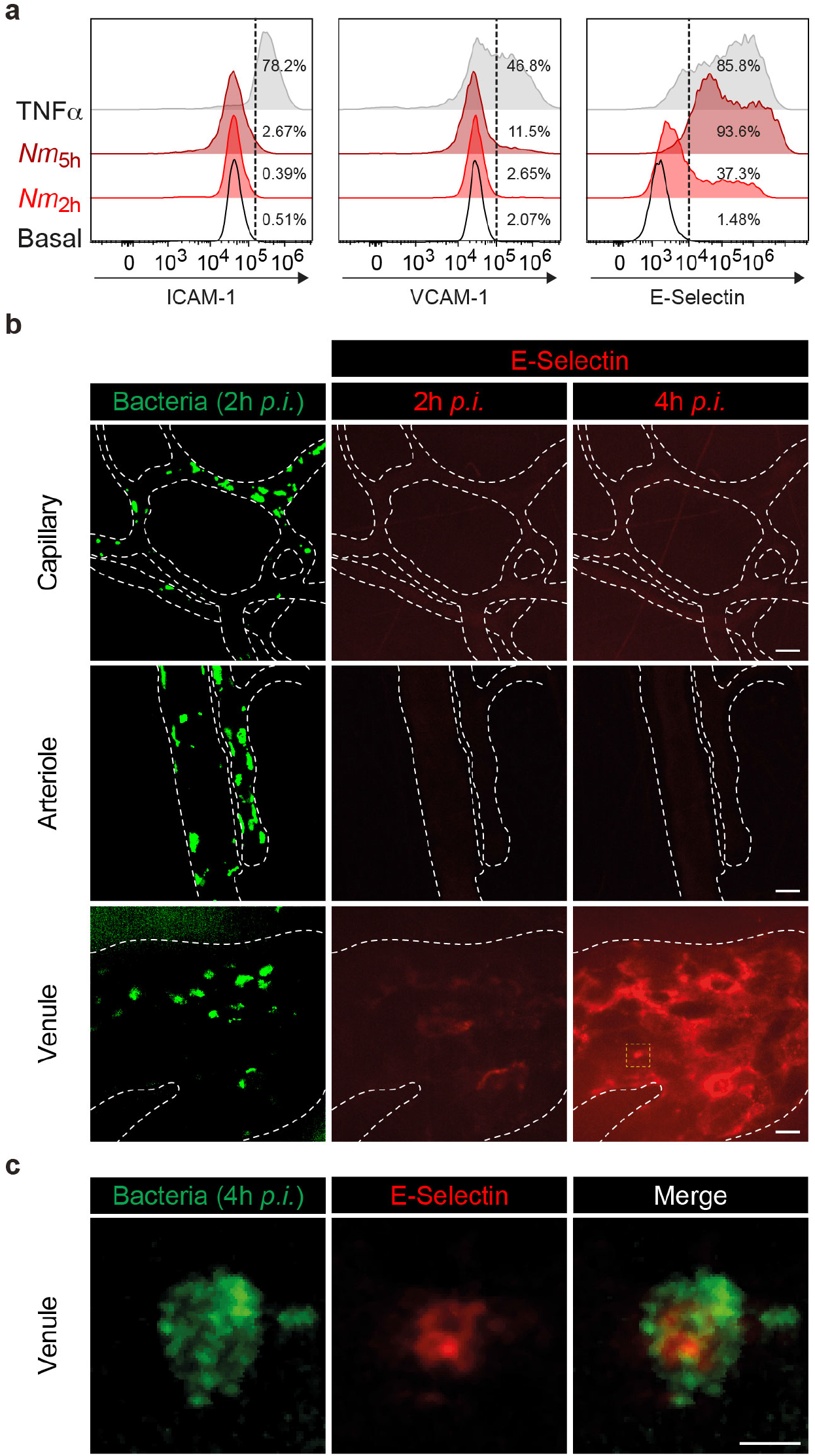
E-Selectin endothelium surface expression is differentially upregulated according to the vascular bed upon infection. **a,** Flow cytometry analysis of cell surface expression of ICAM-1, VCAM-1 and E-Selectin (CD62E) on HUVEC cells under resting conditions (Basal) or following either infection with *Neisseria meningitidis* for 2h (*Nm*_2h_) or 5h (*Nm*_5h_) or an over-night incubation with 20 ng.ml^−1^ TNFα. Data are representatives of N=3 independent experiments. The percentages of positive cells (above the dashed lines) are shown per condition and marker. **b,** *In vivo* expression of E-Selectin at the surface of different human vessel types (Capillary, Arteriole and Venule) following two (2h *p.i.*) and four (4h *p.i.*) hours of infection. Images are representative of N=3 infected mice imaged independently. GFP-expressing *Neisseria meningitidis* appear in green and E-Selectin in red following *in vivo* labelling by *i.v.* injection of PE-labelled anti-CD62E monoclonal antibody. Dashed lines delineate human vessels (UEA-1 lectin). Scale bar, 20 μm. **c,** High-magnification view (yellow dashed square in panel B) of a bacterial microcolony at 4h *p.i.* on the venular endothelium surface and the local upregulation of E-Selectin expression. Scale bar, 5 μm.

## Discussion

To mount a protective innate immune response, the inflammatory cascade needs to trigger the recruitment of numerous neutrophils to the precise site of infection, leading to the efficient clearance of the pathogen. In the case of fulminant infections caused by meningococci, the innate immune response is not successful at keeping the pathogen in check. Experimental results provided here, confirmed by observations in human cases, explain how this is due to the particular proliferation niche of this bacterium. While meningococci have the ability to colonize different vascular beds; capillaries, venules, and arterioles; only infected venules allow efficient neutrophil recruitment.

The ability of bacteria to adhere to the endothelium wall is the starting point for the innate immune response as this allows the concentration of bacteria at a specific focal point rather than their systemic distribution in the blood. In absence of adhesion, neutrophil recruitment is barely detectable. Bacterial accumulation on the endothelial surface is thus a trigger for neutrophil recruitment. The ability of bacteria to auto-aggregate also enhances neutrophil recruitment. This is likely explained by the amplification of bacterial numbers at the site of infection due to the three-dimensional accumulation of bacteria. It could also be envisioned that the formation of aggregates leads to more perturbations in blood flow and subsequently more inflammatory signals. Nevertheless, type IV pili are central players in triggering the inflammatory response by allowing adhesion and auto-aggregation. Bacterial adhesion also shapes the innate immune response by leading to the colonization of different vessel types including capillaries. In a previous study focusing on meningitis and infection of the brain, we had shown a preferential colonization of capillaries in this organ^23^. The current study, based on an *in vivo* model and clinical samples extends these findings to other organs, confirms infection of capillaries and shows that other vascular beds can be infected.

Venules are readily infected and bacterial aggregates bound to the venular endothelial surface can be seen 2-3 hours *p.i.*. In this context, neutrophils are recruited rapidly and in large numbers at similar times. Neutrophil recruitment correlates with the expression of E-Selectin at the surface of the venular endothelium, which promotes neutrophil adhesion and crawling. Upon venular infection, neutrophils are located inside the vascular lumen, actively crawl on the endothelium surface and phagocytose bacteria. Evidence of the disappearance of aggregates from the endothelium surface upon neutrophil local recruitment is clearly seen, thus demonstrating their functional impact in bacterial clearance. Accordingly, depletion of neutrophils leads to more severe infection as measured by global bacterial numbers inside the tissue and evidence of vascular damage. Interestingly, at early time points post-infection, high resolution intravital microscopy of infected venules allowed direct observation of adhering bacteria being engulfed by neutrophils. Considering the intimate type IV pili-mediated adhesion of bacteria to the endothelial surface, allowing bacterial aggregates to resist high mechanical stress^24^, the phagocytosis of such bacterial colonies by neutrophils thus implies the engagement of intense forces that would warrant further investigation.

The scenario in arterioles and capillaries is very different as bacterial elimination is made difficult by several factors. Bacterial proliferation is barely altered by very limited numbers of recruited neutrophils, if any. The absence of endothelial adhesion receptor expression, such as E-Selectin, in these vascular beds upon infection provide an explanation to the reduced levels of intravascular neutrophil recruitment. At later time points post-infection, vessels are entirely full of bacteria, to the point where the few neutrophils on site are trapped in bacterial colonies. The combination of coagulation and occupation of the luminal space by bacteria and congested red blood cells rapidly stops blood flow and consequently neutrophils have a limited access to reach the bacterial colonies. In the case of arterioles, neutrophils can also be found surrounding vessels in the animal model as observed in the human cases, confirming the relevance of these observations^7^. Such neutrophils likely extravasated from nearby venules and migrated through the tissue towards the infected arterioles. Interestingly, these extraluminal neutrophils fail to enter arterioles from the outside and remain within the perivascular area for hours without any evidence of phagocytosis. Although a matter of debate, the ability of neutrophils within tissues to go back inside venules by a process termed *reverse migration* has been described^25^. In the case of meningococcal infections, the inability of these cells to enter vessels or even reach inside infected arterioles could simply be explained by the nature and thickness of the arteriolar vascular wall. The lumen of arterioles and capillaries thus provide a safe haven for the bacterium.

The question remains as to why intraluminal infection of arterioles is insufficient to induce the expression of adhesion receptors despite intense local infection. In arterioles, it is generally accepted that the expression of such adhesion molecules is lower than in venules, leading to lower levels of interaction of neutrophils with the arteriolar endothelium^26^. Nevertheless, in certain instances, the expression of adhesion molecules in arterioles is significantly increased during inflammation. For instance, TNFα treatment results in increased expression of P- and E-Selectin, as well as ICAM-1 and VCAM-1, in cremaster muscle arterioles^27^. Angiotensin II has also been reported to induce the recruitment of neutrophils to arterioles^28^. It nevertheless remains that during *N. meningitidis* infection, despite the impressive accumulation of bacteria within the lumen of arterioles, insufficient levels of adhesion molecules are expressed to provoke the efficient recruitment of neutrophils.

The ability of bacteria to escape the innate immune response and accumulate inside vessels is crucial in triggering the vascular damages that constitute the hallmark of this disease. They include vascular congestions, intravascular coagulation and loss of vessel integrity. The rapid and massive occurrence of these damages lead to decreased tissue oxygenation and finally organ dysfunction. In addition to determining the role of neutrophils in this process, this study provides a detailed description of the kinetics of bacteria-induced vascular damages. At 2-3 h post-infection, about 30% of vessels in the infected tissue show evidence of congestion on histological slides. At these early time points, barely any neutrophils are recruited, suggesting that they have little or no role in the initial stages of vascular damage. At later stages (16-24h), nearly all vessels of the infected tissue become congested providing a striking illustration of how infection causes organ failure. At these same time points, a fraction of vessels loses their integrity and vascular contents are released in the tissue. Our study shows that neutrophils play a protective role in this second phase of the infection by limiting the extent of vascular leakage. This is associated with increased congestion suggesting that neutrophils could enhance coagulation. Exactly how neutrophils limit vascular damage remains to be determined. Assuming that the bacteria themselves trigger the vascular damage, the effect of neutrophils on the number of bacteria could be sufficient to explain their protective role on the vessels. Alternatively, neutrophils could facilitate the coagulation response as previously seen in other models^29^. The specific impact of neutrophils on coagulation during *N. meningitidis* infection needs to be further explored as it could potentially limit further colonization by bacteria.

The results of this study provide an explanation of the fulminant progression of *N. meningitidis* infection by showing how the adhesive and aggregative properties of the bacteria protect them from the function of the innate immune cells. Meningococci occupy the vascular lumen of different vascular beds and most importantly arterioles and capillaries to which neutrophils are poorly recruited following infection. As a consequence, the uncontrolled accumulation of bacteria inside such vessels leads to vascular damages that characterize this fulminant life-threatening infection. This particular means of evading the immune response by targeting a form of immune privileged site could be involved in the high mortality still associated with *purpura fulminans*^30^.

## Methods

### Mice

SCID/Beige (CB17.Cg-*Prkdc*^*scid*^*Lyst*^*bg-J*^/Crl) mice, which were used in all the experiments performed in this study except for the intravital imaging of neutrophils during the late phase of the infection, were purchased from Charles Rivers (France). *Rag*_*2*_ ^−/−^*γ*_*c*_ ^−/−^ mice (kindly provided by the Central Animal Facility, Institut Pasteur, Paris, France) were breeded with *LysM*^*gfp*/+^ mice (kindly provided by Ivo Gomperts Boneca, Institut Pasteur, Paris, France) to obtain *Rag*_*2*_ ^−/−^*γ*_*c*_ ^−/−^*LysM*^*gfp*/+^ mice that were used for the intravital imaging of neutrophils during the late phase of the infection. All mouse strains were housed under specific pathogen-free condition at Institut Pasteur. Mice were kept under standard conditions (light 07.00-19.00h; temperature 22±1°C; humidity 50±10%) and received sterilized rodent feed and water *ad libitum*. All experiments were performed in agreement with guidelines established by the French and European regulations for the care and use of laboratory animals and approved by the Institut Pasteur committee on Animal Welfare (CETEA) under the protocol code CETEA 2015–0025. For all experiments, male and female mice between 6 and 10-weeks of age were used. Littermates were randomly assigned to experimental groups.

### Human skin

Normal human skin was obtained from adult patients (20-60 years old), both males and females, undergoing plastic surgery in the service *de chirurgie reconstructrice et plastique* of Groupe Hospitalier Saint Joseph (Paris, France) or the service *de chirurgie plastique, recontructrice et esthetique* of Hôpital Européen Georges Pompidou (Paris, France). In accordance with the French legislation, patients were informed and did not refuse to participate in the study. All procedures were approved by the local ethical committee *Comité d’Evaluation Ethique de l’INSERM* IRB 00003888 FWA 00005881, Paris, France Opinion: 11-048.

### Xenograft model of infection

5-8 weeks old mice, both males and females, were grafted with human skin as previously described^8^. Briefly, a graft bed of approximately 1–2 cm^2^ was prepared on the flank of anesthetized mice (intraperitoneal injection of ketamine and xylazine at 100 mg.kg^−1^ and 8.5 mg.kg^−1^, respectively) by removing the mouse epithelium and the upper dermis layer. A human skin graft (200 μm thick) comprising the human epidermis and the papillary dermis was immediately placed over the graft bed. Grafts were fixed in place with surgical glue (Vetbond, 3M, USA) and dressings were applied for 2 weeks. Grafted mice were used for experimentation 3-6 weeks post-surgery when the human dermal microvasculature is anastomosed to the mouse circulation without evidence of local inflammation, as previously described^8^. All efforts were made to minimize suffering.

### *Neisseria meningitidis* strains and mouse infection

All *N. meningitidis* strains described in this study were derived from the recently sequenced 8013 serogroup C strain (http://www.genoscope.cns.fr/agc/nemesys)^31^. Mutations in *PilD* and *PilC1* genes have been previously described^31,32^. Wildtype (SB), SA, *pilD* and *pilC1* bacterial strains were genetically modified to constitutively express either the green fluorescent protein (GFP)^8^ or the near-infrared fluorescent protein (iRFP)^12,33^ under the control of the *PilE* gene promoter, as previously described. Strains were streaked from −80°C freezer stock onto GCB agar plates and grown overnight in a moist atmosphere containing 5% CO_2_ at 37°C. For all experiments, bacteria were transferred to liquid cultures in pre-warmed RPMI-1640 medium (Gibco) supplemented with 10% FBS at adjusted OD_600nm_ = 0.05, and incubated with gentle agitation for 2 hours at 37°C in the presence of 5% CO_2_. Bacteria were washed twice in PBS and resuspended to 10^8^ CFU.ml^−1^ in 1x PBS. Prior to infection, mice were injected intraperitoneally with 8 mg of human transferrin (Sigma Aldrich) to promote bacterial growth *in vivo* as previously described^8^. Mice were infected by intravenous injection of 100 μl of the bacterial inoculum (10^7^ CFU total). For intradermal infection, 10^6^ bacteria were resuspended in 50 μl 1x PBS containing 8 mg of human transferrin. No influence of mice sex on bacterial colonization have been observed (data not shown).

### Cell culture and infection

Primary human umbilical endothelial cells (HUVECs) were purchased from Lonza (pooled donors, # C2519A) and cultured in EGM-2 complete medium (Lonza, # CC-3162) without antibiotics. Cells were used between passage 2 and 6. For infection, 2.5×10^5^ HUVECs were plated in wells of 6-well plates and stimulated overnight with 20 ng.ml^−1^ human recombinant TNFα (PeproTech/Tebu Bio, #300-01A-A) or infected at MOI=200 with *Neisseria meningitidis* for 30 min, washed to removed non-adherent bacteria and incubated for an additional 2h or 5h, prior to be harvested and stained for flow cytometry analysis (see below).

### *Ex vivo* colony-forming units (CFU) enumeration

To assess bacteraemia (blood circulating bacteria) in infected animals, 10 μl of blood was sampled before infection, 5 min after infection and at the time of sacrifice. Serial dilutions of blood were plated on GCB agar plates and incubated overnight at 37°C and in a moist atmosphere containing 5% CO_2_. Bacterial counts were expressed in colony-forming units (CFU) per ml of blood.

To assess the extent of vascular colonization by meningococci (adherent bacteria) following mouse sacrifice at indicated times post-infection, tissue biopsies were collected using a sterile dermatological biopsy puncher (approximately 4 mm^2^), weighted and placed in 500 μl 1x PBS. Skin biopsies were dissociated and homogenized using MagNA lyser homogenizer (Roche, France) and serial dilutions of skin homogenates were plated on GCB plates incubated overnight at 37°C and in a moist atmosphere containing 5% CO_2_. Bacterial counts were expressed in colony-forming units (CFU) per mg of skin.

### Cell dissociation from skin samples

Cell suspensions were isolated from human skin xenograft or mouse skin as previously described with some modifications^34^. Briefly, skin biopsies were collected immediately after mouse sacrifice and weighed on a fine balance. Subsequently, skin was cut into small pieces and digested with gentle agitation for 60-90 min at 37°C in CO_2_-independent medium containing 25 mM Hepes, 0.4 mg.ml^−1^ Liberase TL (Sigma Aldrich), 0.04 mg.ml^−1^ DNase I (Sigma Aldrich), and 100 U.ml^−1^ Penicillin/Streptomycin (Gibco). 0.1 μg.ml^−1^ Cytochalasin D was added to the digestion mix when assessing bacterial phagocytosis by macrophages. The resulting single cell-suspension was passed through a 70-μm cell strainer (BD Bioscience) and treated with 1x RBC lysis solution (BioLegend). Cells were again filter using 40-μm cell strainer (BD Bioscience) and counted. Cell viability was determined using trypan blue exclusion.

### Flow Cytometry

#### Dissociated skin

Single-cell suspensions were labelled in aliquots of 10^6^ cells per 100 μl in 1x PBS supplemented with 2% FBS, according to standard protocols^35^. Fc receptors were blocked using the anti-mouse CD16/CD32 (FcBlock clone 2.4G2) monoclonal antibodies (BD Biosciences, #553141, 1/400) and cells were stained for 30 min at 4°C with a combination of the following anti mouse immunophenotyping antibodies: from Biolegend: PacificBlue-conjugated anti-Ly-6C (clone HK1.4, #128014, 1/400), PacificBlue- or PE/Cy7-conjugated anti-Ly-6G/Ly-6C (GR-1) (clone RB6-8C5, #108430 or #108416, 1/400 and 1/200, respectively), PacificBlue-conjugated anti-CD45 (clone 30-F11, #103126, 1/200). From eBioscience: APC-conjugated anti-CD11b (clone M1/70, #17-0112-82, 1/200), PE-conjugated anti-Ly-6C (clone HK1.4, #12-5932-82, 1/400). From BD Biosciences: BUV395-conjugated anti-CD45 (clone 30-F11, #564279, 1/200). Exclusion of non-viable cells has been achieved using eFluor780 Fixable viability dye (eBioscience, #65-0865) according to manufacturer’s instructions. After staining, cells were washed in 1x PBS and fixed for 20 min at 4°C with 4% paraformaldehyde in 1x PBS and washed with 1x PBS supplemented with 2% FBS. Data were acquired using a BD LSR Fortessa^TM^ flow cytometer controlled with the BD FACSDiva software (BD Bioscience). Data analysis was carried out using FlowJo software v10 (Tree Star, Ashland, OR, USA). Numbers of cells were expressed per mg of skin according to the following formula: 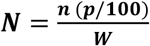 where ***N*** is the number of cells per milligram of skin, ***n*** the number of live cells obtained after skin dissociation (Trypan blue exclusion), ***P*** the percentage of the given cell population among the “live cells” gate determined by flow cytometry and ***W*** the weight in milligram of the corresponding biopsy before skin dissociation.

#### Cultured endothelial cells

Following infection for the indicated times or TNFα treatment, cells were harvested using 37°C preheated Versene (Gibco, #15040066) to preserve cell surface epitopes. Fc receptors were blocked using Human TruStain FcX (BioLegend, #422302, 1/20) diluted in cold FACS buffer (1x PBS supplemented with 0.5% BSA and 2 mM EDTA). Cells were stained for 30 min on ice using AlexaFluor700-conjugated anti-Human CD54/ICAM-1 (clone 1H4, Life Technologies, #MA528553, 1/75), PE-Cy7-conjugated anti-Human CD106/VCAM-1 (clone STA, eBioscience, #25-1069-42, 1/75) and PE-conjugated anti-Human CD62E/E-Selectin (clone P2H3, eBioscience, #12-0627-42, 1/20) diluted in cold FACS buffer. Exclusion of non-viable cells has been achieved using eFluor506 Fixable viability dye (eBioscience, # 65-0866-14) according to manufacturer’s instructions. After staining, cells were washed in cold 1x PBS and fixed over-night at 4°C with 1% paraformaldehyde in 1x PBS. Data were acquired using a CytoFLEX LX controlled with the CytExpert Software (Beckman Coulter) flow cytometer and analysis was carried out using FlowJo software v10 (Tree Star, Ashland, OR, USA).

### Preparation of skin tissue samples for fluorescence microscopy and histology

Skin biopsies were collected from control mice or at different times post-infection, fixed in 4% paraformaldehyde (PFA) for 24h (or 2% PFA up to 15 days) at 4°C, washed at least twice in 1x PBS and dehydrated overnight in 20% (w/v) sucrose at 4°C. Tissue samples were then embedded in OCT (Tissuetek) and frozen at −80°C. 7-10 μm thick cryostat sections were immobilized on Superfrost plus microscopy slides (Thermo scientifics) and prepared either for haematoxylin and eosin (H&E) stain or immunostaining using the Mouse on Mouse kit (Vector laboratories) according to manufacturer’s instructions. Sections were stained using the following antibodies: rat anti-mouse Ly-6C/Ly-6G (RB6-8C5, BD Bioscience, #550291, 1/200) followed by AlexaFluor647-conjugated goat anti-rat IgG (ThermoFisher Scientific, #A21247, 1/200). Human vessels were stained using Rhodamine-conjugated Ulex Europaeus Agglutinin I (UEA-1) lectin (VectorLaboratories, #DL-1067, 1/200). Skin sections were then mounted using Vectashield mounting reagent (Vector laboratories). Fluorescent images were acquired on an inverted spinning-disk confocal microscope (Ti-eclipse, Nikon) equipped with an EMCCD camera (Evolve, Photometrics) and a 40x oil immersion objective using Metamorph Imaging Software (Molecular Devices). Scanning of H&E-stained cryosections was performed using a slide scanner Axio scan Z1 (Zeiss) at 20x magnification and images were processed using ZenBlue software (Zeiss). Image processing was performed using Fiji software^37^ except when mentioned. Images shown in the figures were cropped from large fields, rotated and their contrast and brightness manually adjusted.

### Spinning disk confocal intravital imaging

#### Mouse surgery and Preparation of the skin flap

Intravital imaging of the human xenograft was adapted from^36^. Briefly, 30-minutes prior to surgery, mice were injected subcutaneously with buprenorphine (0.05 mg.kg^−1^) and anesthetized by spontaneous inhalation of isoflurane in 100% oxygen (induction: 4%; maintenance: 1.5% at 0.3 L.min^−1^). A middle dorsal incision was made from the neck to the lower back and the skin supporting the human xenograft was flipped and secure onto an aluminium custom-made heated deck (36°C). The human microvasculature within the graft was exposed by carefully removing the excess of connective tissue. The skin flap was covered with a coverslip maintained thanks to a 3D-printed custom-made holder to avoid any pressure on the xenograft vasculature, sealed with vacuum grease and continuously moistened with warmed 1x PBS (36°C). Mice hydration was maintained by intraperitoneal injection of 200 μl 0.9% saline solution every hour. During the course of the experiment, mouse body temperature was maintained at 37°C using a heating pad and oxygen saturation and heart rate were monitored using the pulse oximetry Physiosuit apparatus (Kent Scientific). The tail vein was cannulated allowing the injection of fluorescent dyes and/or bacteria.

#### Spinning disk confocal microscope

Intravital imaging was performed using a Leica DM6 FS upright microscope equipped with a motorized stage. The microscope is fitted with HCX PL Fluotar 5x/0.15, HC Fluotar 25x/0.95 and HC PL APO 40x/1.10 objectives lens (Leica), mounted on an optical table to minimize vibration and totally enclosed in a biosafety cabinet (Noroit). The microscope is coupled to a Yokogawa CSU-W1 confocal head modified with Borealis technology (Andor). Four laser excitation wavelengths (488, 561, 642 and 730 nm) were used in fast succession and visualized with the appropriate long-pass filters. Fluorescence signals were detected using a sCMOS 2048×2048 pixel camera (Orca Flash v2+, Hamamatsu). Metamorph acquisition software (Molecular devices) was used to drive the confocal microscope.

#### Imaging heterotypic interactions between murine neutrophils and human endothelium

2 hours prior to the skin flap surgery, mice were subcutaneously injected with 0.5 μg of human recombinant TNFα (Preprotech, #300-01A) under the human skin graft. After the skin flap surgery of grafted SCID/beige mice, the murine vasculature was labelled with 15 μg Dylight488-conjugated anti-mouse CD31 (LEAF purified anti-mouse CD31, clone 390, Biolegend, #102412 + DyLight488 Antibody Labelling Kit, ThermoFisher Scientific, #53024) and the human vasculature with 100μg of Dylight755-conjugated UEA-1 lectin (Unconjugated UEA-1 lectin, Vector Laboratories, #L-1060 + DyLight755 Antibody Labelling Kit, Thermofisher Scientific, #84538). Neutrophils were labelled with 2.5 μg Dylight550-conjugated anti-mouse Ly-6G (LEAF purified anti-mouse Ly-6G, clone 1A8, Biolegend, #127620 + DyLight550 Antibody Labelling Kit, ThermoFisher Scientific, #84530) 15 min prior to recordings. Images were recorded 3 and 4 hours after TNFα stimulation. Time-lapse z-stack series (1.5-2 μm spacing) were recorded every 30 seconds for 30 minutes at 3 and 4 hours post TNFα stimulation.

#### Imaging the early phase of infection (0-6h)

After the skin flap surgery of grafted SCID/beige mice and 15 min prior to the tail-vein injection of iRFP-expressing bacteria (10^7^ CFU in 100 μl 1x PBS), 100 μg Dylight755-conjugated UEA-1 lectin (Unconjugated UEA-1 lectin, Vector Laboratories, #L-1060 + DyLight755 Antibody Labelling Kit, Thermofisher Scientific, #84538), 15 μg Dylight488-conjugated anti-mouse CD31 (LEAF purified anti-mouse CD31, clone 390, Biolegend, #102412 + DyLight488 Antibody Labelling Kit, ThermoFisher Scientific, #53024) and 2.5 μg Dylight550-conjugated anti-mouse Ly-6G (LEAF purified anti-mouse Ly-6G, clone 1A8, Biolegend, #127620 + DyLight550 Antibody Labelling Kit, ThermoFisher Scientific, #84530) were injected to label and visualize the human vessels, the mouse dermal microvasculature and the mouse neutrophils, respectively. z-stack images (1.5-2 μm spacing) were captured every 15 minutes for 6 hours.

#### Imaging adhesion of bacteria in arterioles during the early phase of infection (0-3h)

2 hours prior to the skin flap surgery, mice were intravenously injected with Hydrazide AlexaFluor633 (2 mg.kg^− 1^, ThermoFisher Scientific, #30633) to label arteriole walls, as previously described^20^. After the skin flap surgery and 15 min prior to the tail-vein injection of GFP-expressing bacteria (10^7^ CFU in 100 μl 1x PBS), 100 μg Dylight755-conjugated UEA-1 lectin (Unconjugated UEA-1 lectin, Vector Laboratories, #L-1060 + DyLight755 Antibody Labelling Kit, Thermofisher Scientific, #84538) were injected in the mouse tail vein to label and visualize the human vessels. z-stack images (1.5-2 μm spacing) were captured every 30 minutes for 3 hours.

#### Imaging the late phase of infection (>16h)

16 hours prior imaging, grafted *Rag*_*2*_^−/−^*γ*_*c*_^−/−^*LysM*^*gfp*/+^ mice were retro-orbitally injected with iRFP-expressing bacteria (10^7^ CFU in 100 μl 1x PBS). The skin flap surgery was initiated 15 hours post-infection and time-lapse z-stack series (1.5-2 μm spacing) of iRFP-expressing bacteria and eGFP-expressing neutrophils were recorded every 30 seconds for 30 minutes.

#### Imaging human E-Selectin (CD62E) expression during the early phase of infection

After the skin flap surgery and 15 min prior to the tail-vein injection of GFP-expressing bacteria (10^7^ CFU in 100 μl 1x PBS) and PE-conjugated anti-human CD62E (6 μg, Thermofisher Scientific, #12-0627-42), 100 μg Dylight755-conjugated UEA-1 lectin (Unconjugated UEA-1 lectin, Vector Laboratories, #L-1060 + DyLight755 Antibody Labelling Kit, Thermofisher Scientific, #84538) and 50 μg of Hydrazide AlexaFluor633 were injected in the mouse tail vein to label and visualize the human vessels and the arteriole walls, respectively. z-stack images (2 μm spacing) were captured every 60 minutes for 4 hours.

#### Image processing

Image were exported from the Metamorph acquisition software (Molecular Devices) as .tiff images, deconvoluted using Huygens software (Scientific Volume Imaging) and edited using Fiji. Videos were exported as .avi files and edited on Final Cut Pro (Apple). 3D-rendering were generated using Imaris software (Bitplane).

### *In vivo* neutrophil depletion

Neutrophils were depleted in grafted mice by intravenous injection of either 50 μg of anti-mouse Ly-6G/Ly-6C (GR-1) monoclonal antibody clone RB6-8C5 (eBioscience, #16-5931-85) or 100 μg of anti-mouse Ly-6G monoclonal antibody clone 1A8 (eBioscience, #16-9668-85) 24 hours prior to infection. Control animals were injected with the respective rat IgG2β,κ or eBrD isotype control (eBioscience, #16-4031-85 and #14-4321-85, respectively). Neutrophil depletion was confirmed by differential leukocyte count and flow cytometry analysis.

### Evaluation of vascular permeability with Evans blue

Twenty-four hours post-infection, anesthetized mice received an intravenous injection of 100 μl sterile Evans blue dye (0.5% w/v in 1x PBS) that was allowed to circulate for 10 minutes. Mice were then perfused with 5 ml of heparin-containing 1x PBS (2 units.ml^−1^) and 4 mm^2^ biopsies of the human xenografts were harvested using dermatological biopsy punchers. Skin biopsies were weighed, disposed into 1.5 ml tubes containing 300 μl formamide and homogenized using MagNA lyser homogenizer set at 6,000 g for 30 sec. Samples were incubated at 37°C during 60 hours to extract Evans blue dye from the tissues and centrifuged to pellet any remaining tissue fragments. The optical densities at 620 nm of the supernatants were measured using a spectrophotometer and the Evans blue concentration (ng.ml^−1^) of each sample was determined.

### Quantification of vascular damage

Images of H&E stained skin sections were manually analysed and more than 200 blood dermal vessels were assessed for every condition at indicated times and classified as healthy, congested (*i.e.* luminal accumulation of red blood cells) or breached (*i.e.* presence of perivascular red blood cells) vessels. The percentage of vessels in each condition was then determined.

### Statistics

All graphs and statistical analyses were performed with GraphPad Prism 8 (GraphPad Software). No statistical method was used to predetermine sample size. Kolmogorov–Smirnov test was used to assess the normality of all data sets. Scatter dot plots show the Mean ± SEM. P-values were considered as statistically significant when inferior at 0.05 with *p<0.05; **p<0.005; ***p<0.0005; ****p<0.0001 and ns, not significant. Statistical details of experiments (sample size, replicate number, statistical significance) can be found in the figures and figure legends.

## Data availability

Data supporting this work are available in the paper. Further information and materials related to the findings of this study are available from the corresponding authors upon request.

## Acknowledgements

The authors thank Elisa Gomez Perdiguero, Molly Ingersoll, Ana-Maria Lennon-Duménil and Benoît Marteyn for fruitful discussions and critical reading of the manuscript. We also thank Albane Imbert (FabLab Institut Pasteur (FLIP), DRTE, Institut Pasteur, Paris, France) for 3D-printing, Michael Hivelin (Hôpital Européen Georges Pompidou) for providing skin samples, Sophie Novault (Cytometry and Biomarkers UTechS, Institut Pasteur, Paris, France) for technical advices, Eric Camerer (UMR-970, Paris-Cardiovascular Research Center PARCC, Paris, France) for technical advices on vascular permeability experiments and the members of the UtechS Photonic BioImaging (Imagopole, C2RT, Institut Pasteur, Paris, France), supported by the French National Research Agency (France BioImaging; ANR-10– INSB–04; Investments for the Future) for assistance in image acquisition and analysis, Patrick Bruneval (UMR-970, PARCC) for the analysis of human cases pathology. This work was supported by the Integrative Biology of Emerging Infectious Diseases (IBEID) laboratory of excellence (ANR-10-LABX-62), and the VIP European Research Council-starting grant (310790-VIP, G.D.). D.O. was supported by a Pasteur-Roux Postdoctoral Fellowship from the Institut Pasteur. H.E.-R. was supported by a Marie Curie Intra-European Fellowship within the 7th project European Community Framework Program (624740).

## Author contributions

G.D. and V.M. designed research; V.M., P.N., T.U., H.E.-R., K.M., P.F. and D.O. performed experiments; T.S. provided human skin samples; V.M., P.N. and D.O. analysed data; D.O. assembled figures; D.O. and V.M. participated in manuscript writing and G.D. supervised overall data analysis and wrote the manuscript.

---

**Supplementary Figure 1.**
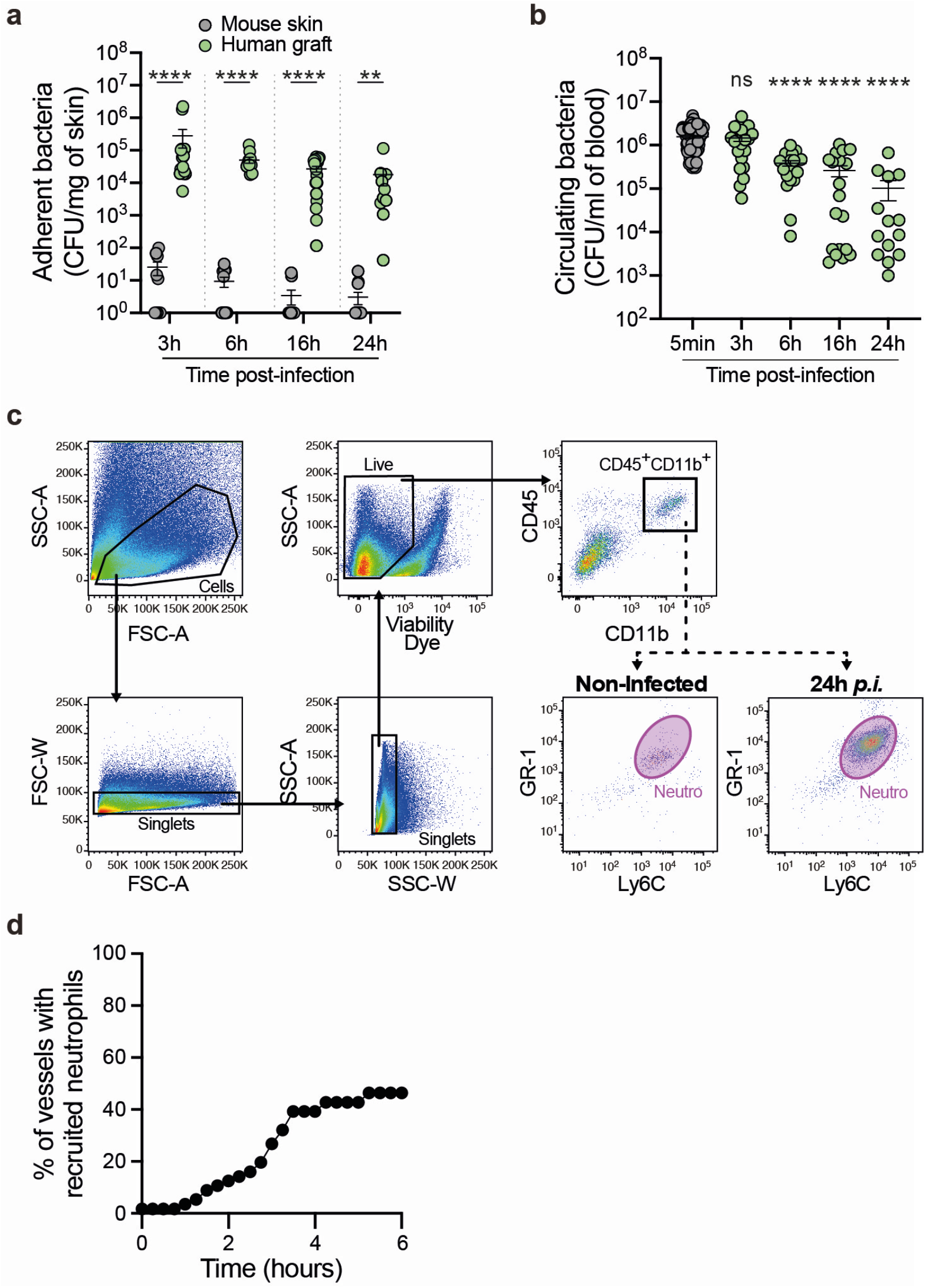
Neutrophil recruitment upon vascular colonization. **a-b,** Bacterial colony forming unit (CFU) counts from **(a)** dissociated skin biopsies (adherent bacteria) collected from either the human grafted or contralateral mouse skin and **(b)** blood (circulating bacteria) of infected mice at the indicated times post-infection. Kruskal-Wallis test with Dunn’s correction for multiple comparisons. n≥10 mice per time point, in total, pooled from N≥4 independent experiments per time point. **c,** Flow cytometry gating strategy used to identify murine neutrophils (Neutro) in dissociated human xenografts based on the cell surface expression of CD45, CD11b, Ly-6C, and GR-1. **d,** Videos obtained from intravital imaging were used to quantify the percentage of vessels effectively recruiting neutrophils during the first 6 hours of the infection. Data are shown as percentage of vessels. Quantifications were performed on n=56 vessels, in total, pooled from N=7 infected mice imaged independently. ns, not significant; *p<0.05; **p<0.005; ***p<0.0005 and ****p<0.0001.

**Supplementary Figure 2.**
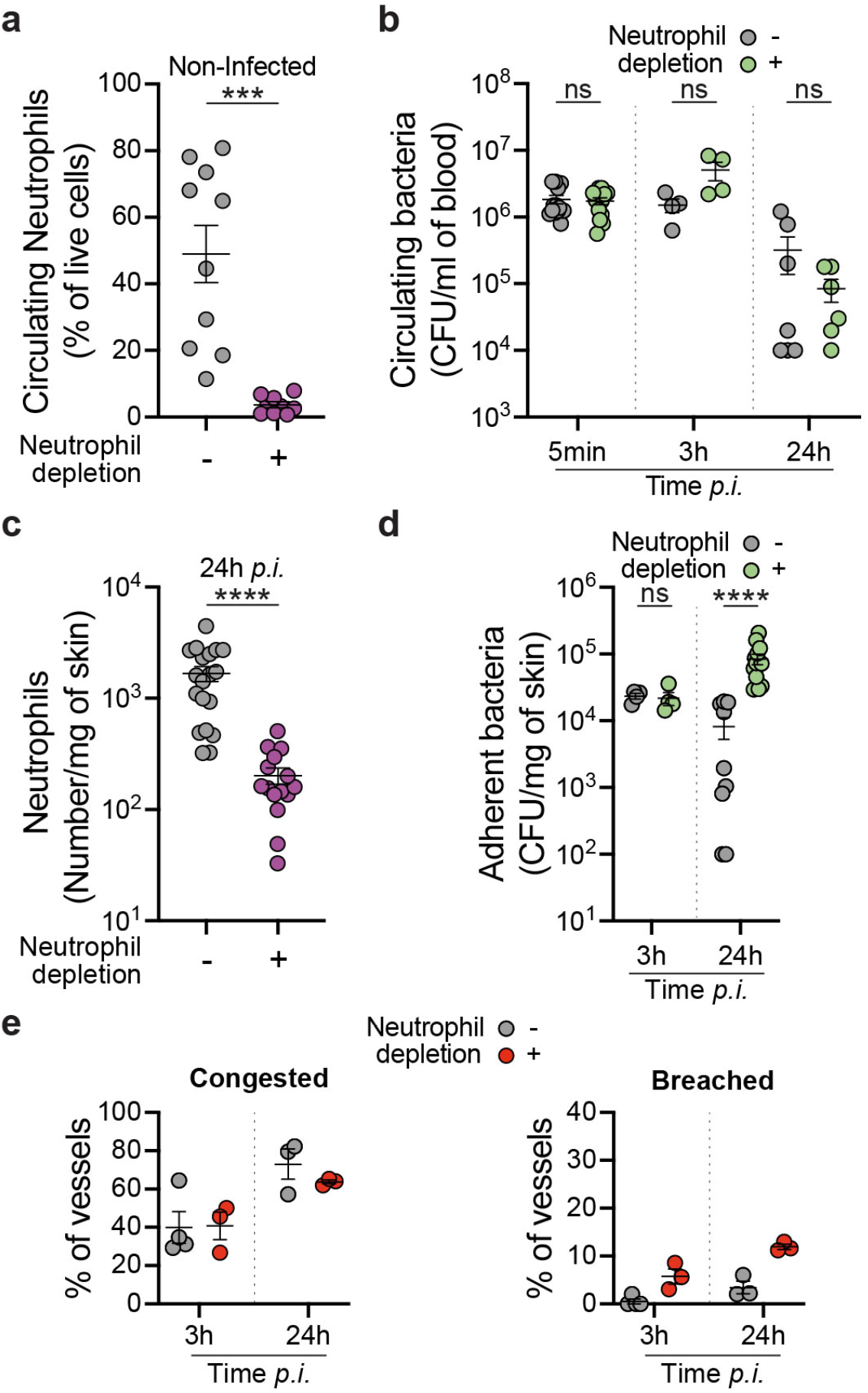
Neutrophils limit both vascular colonization by meningococci and vessel damages. **a-d,** Neutrophil depletion was achieved by intravenous injection of the neutrophil-depleting antibody (anti-Ly-6G, clone 1A8) 24h prior to mouse infection. Mice were sacrificed 24h post-infection and analyses were carried out. **a,** Percentage of blood circulating neutrophils in non-infected mice pre-treated with either the isotype control antibody (−) or the neutrophil-depleting antibody (+). Two-tailed Unpaired t test. n≥9 mice per group, in total, pooled from N=4 independent experiments. **b,** Bacterial colony forming unit (CFU) counts from blood (circulating bacteria) of mice pre-treated with either the isotype control antibody (−) or the neutrophil-depleting antibody (+) and infected for the indicated times. Kruskal-Wallis test with Dunn’s correction for multiple comparisons. n≥4 mice per group, in total, pooled from N=3 independent experiments. **c,** Neutrophil numbers in human xenografts of mice pre-treated with either the isotype control antibody (−) or the neutrophil-depleting antibody (+) and infected for 24 hours. Two-tailed Man-Whitney test. n≥15 mice per group, in total, pooled from N=5 independent experiments. **d,** Bacterial colony forming unit (CFU) counts from dissociated human xenografts (adherent bacteria) collected from mice pre-treated with either the isotype control antibody (−) or the neutrophil-depleting antibody (+) and infected for the indicated times. Kruskal-Wallis test with Dunn’s correction for multiple comparisons. n≥4 mice per group, in total, pooled from N=2 (3h) and 3 (24h) independent experiments. **e,** Quantification of vascular damage upon mouse infection for the indicated time points following neutrophil depletion by intravenous injection of the neutrophil-specific depletion antibody (anti-Ly-6G, clone 1A8) 24h prior to mouse infection. Quantification were performed on n≥200 vessels, in total, pooled from N=3 mice per time point. ns, not significant; *p<0.05; **p<0.005; ***p<0.0005 and ****p<0.0001.

**Supplementary Figure 3.**
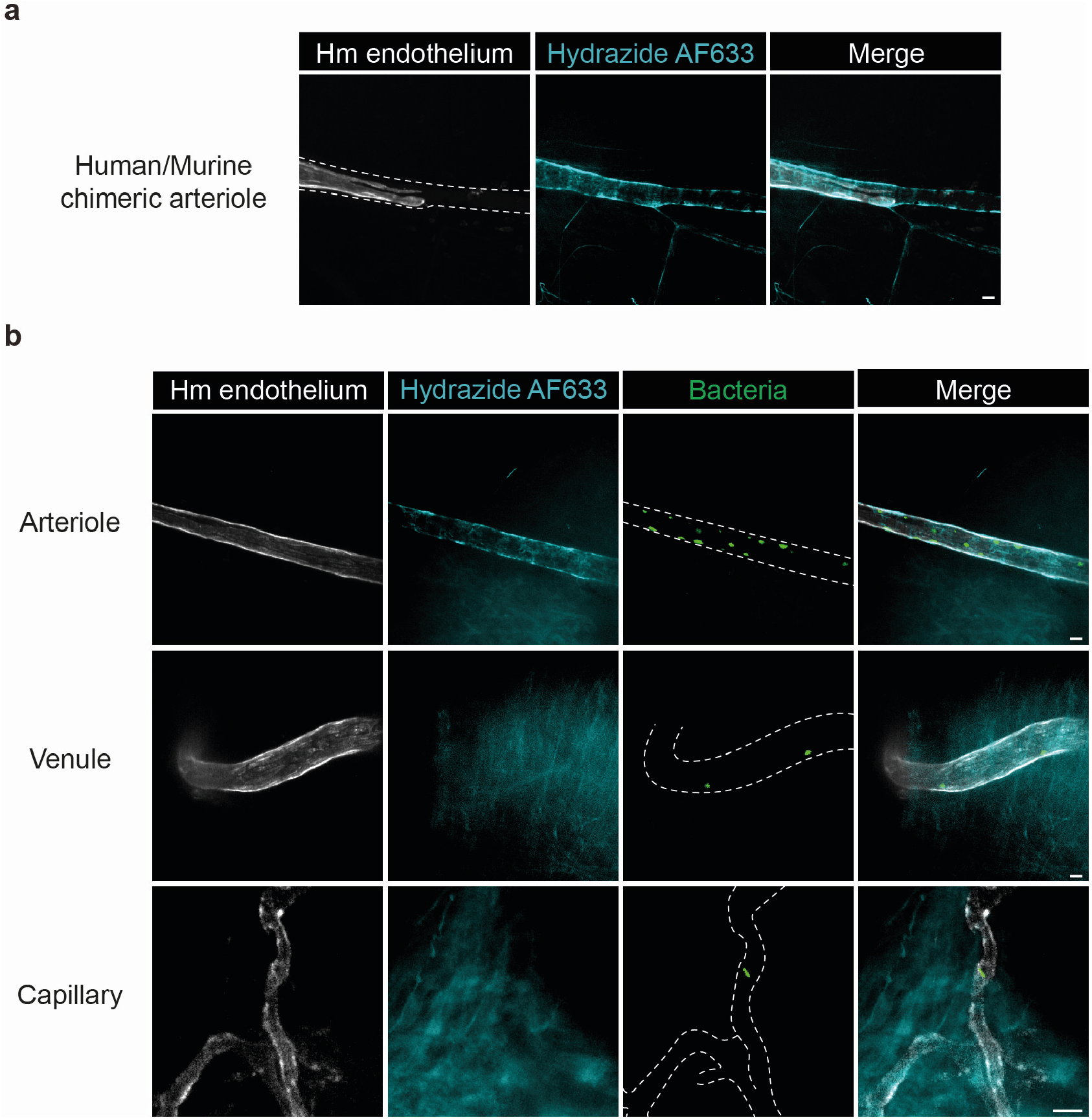
Identification of the different vessel types based on hydrazide AlexaFluor633 labelling upon infection. **a,** Representative images of vessel labelling using Hydrazide AlexaFluor633 (AF633) in a chimeric human/murine arteriole, confirming that Hydrazide AF633 equally stains human and murine arterioles. **b,** Representative images of vascular colonization of the different human vascular beds 1h post-infection by GFP-expressing *Neisseria meningitidis* (green) as revealed by the UEA-1 lectin (human endothelium, grey) and Hydrazide AF633 (arterioles, cyan) double labelling. Scale bar, 20 μm.

## Supplementary Video Legends

**Supplementary Video 1. Vascular colonization by *Neisseria meningitidis.*** Intravital imaging of iRFP-expressing *Neisseria meningitidis* vascular colonization. Bacteria rapidly bind to the human endothelium (UEA-1 lectin, grey) and locally proliferate. At 6 hours post-infection, the 3D-rendering shows the complete colonization of the human endothelium whereas no bacteria were detected within adjacent mouse vessels (mouse CD31, red). Time, hh:min:sec. Scale bar, 50 μm.

**Supplementary Video 2. Heterotypic mouse-human interactions.** Intravital visualization of the interaction between mouse neutrophils (Ly-6G, magenta) and TNFα-mediated inflamed human (UEA-1 lectin, grey) and/or mouse (mouse CD31, red) endothelia. A chimeric human/mouse vessel is shown in the first field of view. Time, hh:min:sec. Scale bar, 20 μm. A segment of a human vessel is shown in the second field of view. Time, hh:min:sec. Scale bar, 10 μm.

**Supplementary Video 3. Neutrophil recruitment during the late phase of *Neisseria meningitidis* infection.** Intravital imaging of neutrophil (*LysM*^*gfp*/+^, magenta) recruitment 16h post-infection with iRFP-expressing *Nm* (green). Neutrophils were massively recruited around the infected human vessel. Tracking of individual neutrophils showed their directed migration towards the infected human vessel. Time, hh:min:sec. Scale bar, 50 μm.

**Supplementary Video 4. Crawling of neutrophils on the human venular endothelium and phagocytosis of adherent *Neisseria meningitidis*.** Intravital visualization of neutrophils (Ly-6G, magenta) internalizing adherent iRFP-expressing *Neisseria meningitidis* (green) inside infected human venules (UEA-1 lectin, grey). In the first field of view, imaging was performed at 3h00 post-infection. Time, hh:min:sec. Scale bar, 20 μm.

**Supplementary Video 5. Perivascular neutrophil recruitment during the early phase of arteriolar colonization by *Neisseria meningitidis.*** Intravital visualization of neutrophil (Ly-6G, magenta) dynamics in the perivascular area of an infected human arteriole (UEA-1 lectin, grey) during the first 6h of infection. Neutrophils remain outside of vessels and do not internalize iRFP-expressing bacteria (green).

**Supplementary Video 6. Reduced neutrophil dynamics following vascular colonization of human arterioles.** Intravital visualization of neutrophil (Ly-6G, magenta) dynamics in an infected human arteriole (UEA-1 lectin, grey) 7h00 post-infection. Neutrophils containing engulfed iRFP-expressing bacteria (green) display a very low motility compared to neutrophils present in the adjacent non-infected mouse venules (mouse CD31, red). Time, hh:min:sec. Scale bar, 20 μm.

